# Pyruvate-driven hydrogen production promotes polyphenol bioconversion by gut bacteria

**DOI:** 10.64898/2026.04.18.719167

**Authors:** Mohamad Eshaghi Gorji, Pui-Kei Lee, Jianing Liu, Linggang Zheng, Xiong Xia, Yu Xinyu, Meng Ziyi, Molly Meng-Jung Li, Lei Dai, Danyue Zhao

## Abstract

Gut microbial biotransformation of poorly absorbable polyphenols into bioactive, bioavailable metabolites is increasingly recognized as a key mechanism underlying their health benefits of polyphenols. Microbial ellagic acid (EA)-to-urolithin conversion represents a typical example, but the environmental factors that facilitate such metabolism remain underexplored. We discovered that urolithin production by a gut commensal bacterium, *Gordonibacter urolithinfaciens* (*G. uro*), is metabolically repressed by arginine. To overcome such limitations, we developed PhenolBoost Medium (PBM) that induces a metabolic shift by suppressing the arginine deiminase pathway while activating pyruvate metabolism and hydrogen production in *G. uro*, thereby driving urolithin dehydroxylation. Transcriptomic profiling and ^13^C-isotopic tracing analysis revealed that pyruvate metabolism in PBM upregulates hydrogenase expression, facilitating the dehydroxylation of EA. PBM also promoted the complete conversion of EA to urolithin A in *G. uro*-*Enterocloster bolteae* co-culture, and other polyphenol biotransformations. In addition, co-culturing *G. uro* with hydrogen-producing *Bacteroides* species significantly increased urolithin production. Furthermore, an arginine-limited, pyruvate-enriched dietary regimen proved effective *in vivo*, resulting in significantly higher urolithin production and bioavailability in a mouse model. Our findings reveal the critical role of hydrogen in facilitating polyphenol dehydroxylation, and offer a viable nutritional strategy for boosting microbial production of beneficial metabolites from polyphenols.

## INTRODUCTION

Polyphenols are plant secondary metabolites widely found in human diets with diverse purported health benefits ^1, 2^. Frequent consumption of polyphenols, naturally abundant in plant-based foods and beverages, has been closely associated with positive health outcomes, including improved cardiometabolic health, reduced oxidative stress and risks of major chronic diseases ^3, 4^. Ellagic acid (EA), naturally abundant in berries, pomegranates and nuts, is one of the most consumed dietary polyphenols with multifarious health benefits ^5, 6^. EA is present either in free form (a phenolic acid) or conjugated as ellagitannins (hydrolysable tannins), but is barely absorbed in the upper intestine and reaches the colon mostly intact. Consequently, its health benefits are largely attributed to the metabolic activity of gut bacteria ^7, 8^.

The catabolism of EA involves a series of metabolic steps that yield urolithin metabolites, hydroxylated 6H-dibenzo[b,d]pyran-6-one derivatives with diverse hydroxylation patterns. Among the urolithins widely detected in humans and animal models, urolithin C (UroC, the key rate-limiting intermediate) and urolithin A (UroA, the most prevalent and bioactive end-product), have received substantial research interest due to their high bioavailability and positive association with favorable metabolic outcomes ^9, 10, 11^. This metabolic pathway starts with the decarboxylation of EA to yield urolithin M5 (UroM5), followed by stepwise dehydroxylations to produce UroC, and then further dehydroxylated to UroA by gut commensals, such as *Enterocloster bolteae* ^12, 13^.

Despite extensive research on EA and urolithins, only a few urolithin-producing gut bacteria have been identified in humans, predominantly belonging to the *Eggerthellaceae* family, notably *Gordonibacter urolithinfaciens* (*G. uro*) and *G. pamelaeae* (*G. pam*). Recent studies have identified the enzyme repertoire responsible for EA transformation by *G. uro* and *G. pam*. Dong et al. (2025) established *G. uro* as a versatile heterologous host for expressing and characterizing these enzymes, successfully mapping multiple previously unknown polyphenol-metabolizing pathways ^14^. Subsequently, Bae et al. (2026) identified members of two distinct molybdenum-dependent enzyme families within *Gordonibacter* spp., i.e., the DMSO reductase and xanthine oxidase families, that function as regioselective catechol dehydroxylases ^15^. These new discoveries have significantly advanced our understanding of EA-to-urolithin conversion.

While *in vivo* conversion of EA to urolithins is generally efficient ^16, 17^, including in our animal study (**Supplementary Fig. 1**), this process exhibits considerable inter-individual variability, leading to ‘high-producer’ and ‘low-producer’ metabotypes in humans ^18, 19^. This robust *in vivo* yield, however, stands in sharp contrast to the poor efficiency of this transformation under current *in vitro* laboratory models. More critically, when urolithin-producing bacteria such as *G. uro* are cultured using standard growth media, the efficiency of initial metabolic steps, i.e., from EA to UroC, is remarkably low. To date, little research has been conducted to identify the key factors that influence these catabolic activities in corresponding gut bacteria, leaving the environmental and nutritional triggers of urolithin production largely unknown.

In this study, we integrate transcriptomics, metabolomics, stable isotope tracer analysis, and *in vivo* studies to uncover the environmental factors governing EA conversion by gut bacteria such as *Gordonibacter* spp. To overcome the limitations of EA-to-urolithin conversion *in vitro*, we developed a novel medium, PhenolBoost Medium (PBM), that induced an “arginine-to-pyruvate” metabolic shift in *G. uro*, resulting in significantly promoted EA-to-urolithin conversion. This shift promoted hydrogen production in the bacteria, serving as the primary driver for enhanced polyphenol dehydroxylation. This mechanism has also been successfully confirmed in multiple instances of catabolism of polyphenols containing pyrogallol and catechol moieties by corresponding bacteria. Furthermore, we demonstrated that this arginine-to-pyruvate shift also significantly increased urolithin production in mice. Together, our study provides novel insight into urolithin metabolism, identifies hydrogen as a key regulator of polyphenol dehydroxylation, and offers a viable nutritional strategy for potentiating microbial catabolic activities which may be applied in industrial biosynthesis of bioactive phenolic metabolites.

## RESULTS

### Standard growth media limit EA-to-urolithin conversion by *G. uro in vitro*

While nutrient-rich media are widely utilized to recover the metabolic potential of anaerobic bacteria isolated from the mammal gut, their efficacy in supporting xenobiotic biotransformation involving multi-step dehydroxylation, such as those in EA-to-urolithins, has not been assessed. Thus, we first sought to establish a benchmark using three standard growth media for anaerobic bacteria including BHI+, PYG+, and WCM. While all three provide the essential nutrients required to support the growth of *G. uro* ^12, 15^, BHI+ and PYG+ are supplemented with additional arginine, and WCM only contains the naturally-present low amount (<1 g/L). In addition, they differ in carbohydrate composition (PYG+ and BHI+ are glucose-rich; WCM is glucose-free but pyruvate-rich). Furthermore, WCM and PYG+ are enriched with hemin, while BHI+ lacks this growth factor (**Supplementary Table 2**).

Despite the growth-promoting capacity of the tested media, none of them was effective in supporting the conversion of EA to UroC (**Fig. 1B**). For *G. uro*, EA-to-UroC conversion in BHI+ and PYG+ was negligible (<1%), with total urolithin yields less than 17%. WCM performed slightly better, giving a total EA degradation of 22.7 ± 1.2%, but UroC production remained extremely low (1.6 ± 0.1%). Similarly, *G. pam* exhibited less than 1% UroC production in all media, corroborating the ineffectiveness of these media in supporting EA biotransformation by *Gordonibacter* spp.

**Fig. 1.**
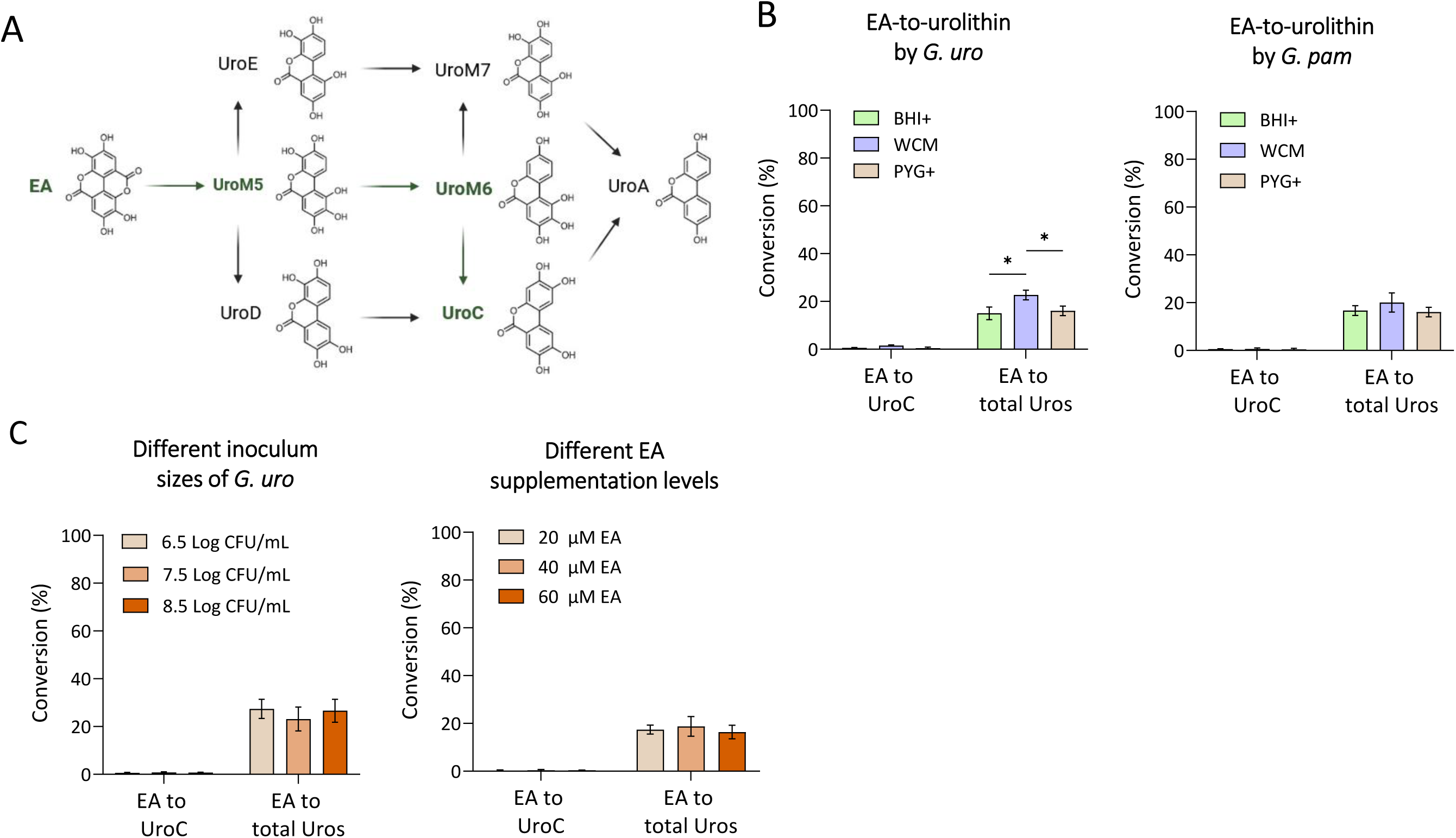
Standard growth media limit *Gordonibacter*-initiated EA-to-urolithin conversion. **(A)** Schematic representation of the microbial metabolic pathway from EA to urolithin intermediates. *G. uro* sequentially converts EA to UroM5, UroM6, and finally UroC. **(B)** EA conversion to urolithin C (UroC) and total urolithins (Uros) in BHI+, WCM, and PYG+ media by *G. uro* and *G. pam*. Bacterial cultures were supplemented with EA (40 µM) and incubated anaerobically for three days. **(C)** EA conversion to UroC and total Uros in PYG+ with different inoculum sizes and EA supplementation levels. All data are presented as mean ± SEM (*n* = 3). Statistical significance was assessed using one-way ANOVA followed by Tukey’s post hoc test (*, P < 0.05; **, P < 0.01; ***, P < 0.001).

We next sought to determine whether such low conversion rates were due to suboptimal culture conditions. To this end, PYG+ was selected as the representative condition for examining inoculum size and substrate concentration, given that it supported the highest bacterial biomass among the three media (**Supplementary Fig. 2**) and was used as the basis for subsequent experiments. We found that the initial inoculum size (6.5, 7.5, and 8.5 log CFU/mL) did not exert any significant effects on EA-to-urolithin conversion (**Fig. 1C**), with UroC conversion consistently below 0.8% and total urolithin conversion below 28%. Similarly, changing the initial EA concentration (20, 40, and 60 µM) yielded no significant improvement (**Fig. 1C**). These results suggest that the metabolic bottleneck for UroC production in these standard media cannot be overcome by simply increasing bacterial inoculum or substrate availability.

### Arginine-to-pyruvate nutritional shift unlocks EA-to-urolithin converting capacity in *Gordonibacter*

A previous metabolomic study in *E. lenta,* a species within the *Eggerthellaceae* family and phylogenetically close to *Gordonibacter* spp., demonstrated that arginine metabolism is the primary source of ATP and drives sustained bacterial growth ^20^. Meanwhile, despite the presence of glucose in the PYG+ medium, *G. uro* is unable to metabolize it ^21^. Therefore, we considered supplementing the medium with pyruvate as an alternative energy source. This decision was supported by two observations: first, pyruvate is a key component of WCM, which is routinely used to cultivate fastidious gut bacteria; and second, our genomic analysis (NCBI database) indicated that *G. uro* harbors the genes necessary for pyruvate utilization. Therefore, we hypothesized that pyruvate supplementation combined with arginine limitation would induce a metabolic reprogramming in *G. uro*. To test this, we evaluated EA biotransformation in four distinct medium conditions: PYG+, arginine-free PYG+ (PYG+(-arg)), PYG+ supplemented with pyruvate (PYG+(+pyr)), and PBM.

The removal of arginine (PYG+(-arg)) led to a modest increase in biotransformation rates, yielding 3.1 ± 1.2% UroC and 22.9 ± 1.5% total urolithins, suggesting that arginine restriction partially relieves EA metabolic repression (**Fig. 2A**). Furthermore, PBM, in which arginine was depleted and pyruvate was supplemented, resulted in a dramatic metabolic activation, achieving a UroC conversion of 28.4 ± 1.5% and a total urolithin yield of 83.0 ± 2.2%. A similar trend was observed in *G. pam*, where PBM increased UroC conversion to 30.8 ± 2.4%. Of note, supplementing pyruvate to the arginine-rich PYG+ medium failed to replicate such benefits with EA-to-UroC conversion rates below 0.2%, confirming that arginine repression dominates over pyruvate activation.

**Fig. 2.**
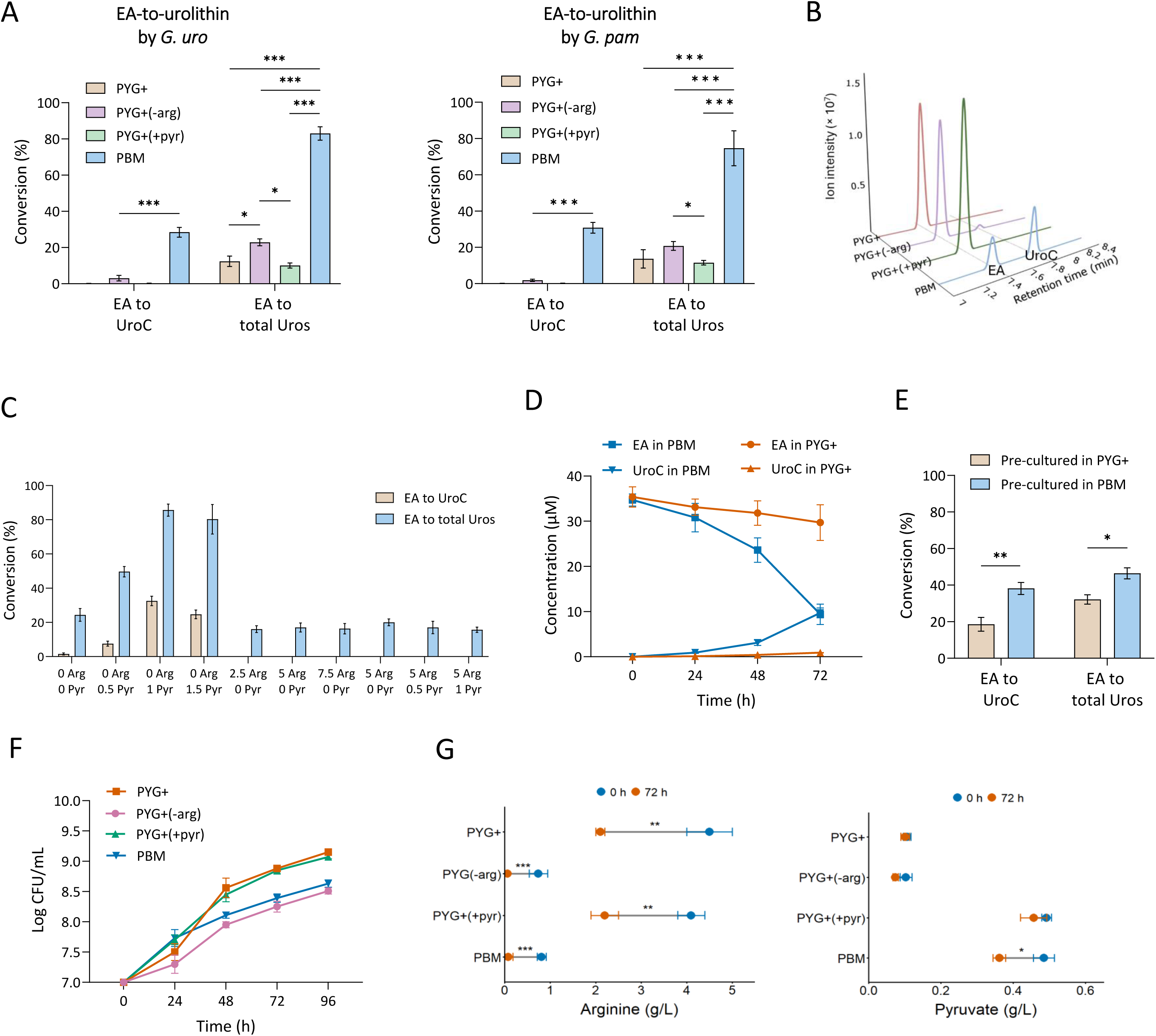
PhenolBoost Medium (PBM) enhances EA-to-urolithin conversion by *Gordonibacter* spp. **(A)** Conversion of EA to UroC and total Uros in PYG+, PYG+(-arg), PYG+(+pyr), and PBM by *G. uro* and *G. pam.* Bacterial cultures were supplemented with EA (40 µM) and incubated anaerobically for 72 h. **(B)** Representative LC-MS chromatograms showing EA and urolithin profiles in the four media. **(C)** Effect of varying concentrations of arginine and pyruvate on EA-to-Uros conversion. **(D)** Kinetics of EA metabolism and UroC production in PYG+ and PBM. **(E)** EA-to-Uros conversion by *G. uro* lysates. *G. uro* was pre-cultured with EA in PYG+ or PBM followed by incubating cell lysate with EA for 24 h. **(F)** Growth of *G. uro* in PYG+, PYG+(-arg), PYG+(+pyr), and PBM. (**G)** Arginine and pyruvate concentrations before (0 h) and after (72 h) *G. uro* growth in PYG+, PYG+(-arg), PYG+(+pyr), and PBM media. All data are presented as mean ± SEM (*n* = 3). Statistical significance was assessed using one-way ANOVA followed by Tukey’s post hoc test (multiple groups) or Student’s t-test (two groups) (*, P < 0.05; **, P < 0.01; ***, P < 0.001).

To determine the optimal medium conditions for EA catabolism, we assessed the effects of varying compositions of arginine and pyruvate. Consistently, EA-to-urolithin transformation was inhibited by arginine: supplementation as low as 2.5 g/L prevented UroC production, regardless of pyruvate levels (**Fig. 2C**). Conversely, in the absence of arginine, pyruvate supplementation enhanced conversion in a dose-dependent manner, peaking at 1 g/L pyruvate.

Time-course analysis of EA and urolithins in PBM demonstrated that UroC production was concurrent with gradual EA depletion (**Fig. 2D**). To verify that such enhanced activity was due to the induction of intracellular enzyme proteins, we incubated EA with soluble protein fraction of the cell lysates. Lysates derived from PBM-cultured cells exhibited significantly higher UroC production efficiency (38.2 ± 2.2%) compared to PYG+-cultured lysates (18.6 ± 3.1%) (**Fig. 2E**). Growth kinetics confirmed that PBM, while yielding lower stationary phase biomass than PYG+ (8.63 vs 9.15 log CFU/mL), created an arginine deficiency (<0.1 g/L within 72 h) that favors catabolic enzyme expression in *G. uro* (**Fig. 2F**).

To assess the utilization of the two supplemented components by *G. uro*, we quantified arginine and pyruvate levels during fermentation. In arginine-supplemented media (PYG+ and PYG+(+pyr)), arginine was significantly consumed by 72 h, with 46.6% and 45.1% left on average, respectively (**Fig. 2G**). In the arginine-deficient media (PBM and PBM(-pyr)), the initial concentration of arginine was below 0.8 g/L, possibly originating from the protein components, and was subsequently depleted after the 72-h incubation. In the meantime, analysis of pyruvate levels suggests that its active consumption requires the absence of arginine (**Fig. 2G**). Pyruvate concentrations significantly decreased by 24.6% in PBM, and by 28.3% in PYG+(-arg) over 72 hours (P < 0.05). In contrast, pyruvate reduction was not statistically significant in both media supplemented with arginine (PYG+ and PYG+(+pyr)), confirming that the presence of arginine represses pyruvate utilization. These findings underscore that while arginine drives robust biomass accumulation, its restriction is essential to the activation of the specific metabolic machinery for urolithin production in *G. uro*. This growth-focused strategy in arginine-rich conditions is further reflected by the high levels of isovaleric acid (**Supplementary Fig. 3**)—likely generated via the overflow metabolism of leucine—whereas this metabolite was absent in PBM and PYG+(-arg). Consistently, higher extracellular levels of the leucine-containing tripeptide Leu-Gly-Pro were also detected in PYG+ compared to PBM (**Fig. 3B**), further supporting the distinct metabolic states between the two conditions.

**Fig. 3.**
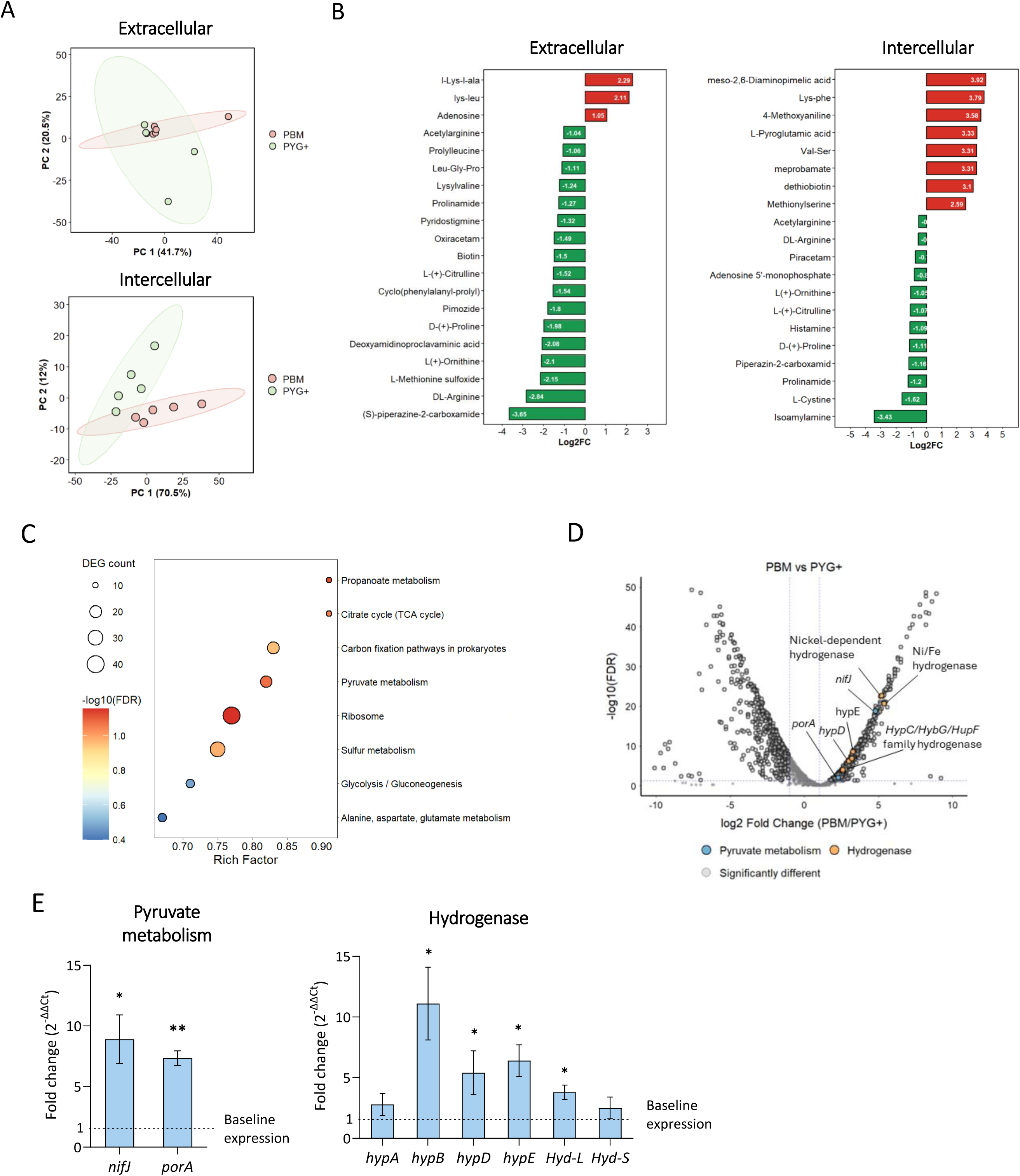
Metabolomic and transcriptomic profiling reveal metabolic shifts in *G. uro* from arginine to pyruvate utilization in PBM. **(A)** PCA plots showing extracellular (top) and intercellular (bottom) metabolite profiles of *G. uro* cultured in PYG+ and PBM. **(B)** Top 20 extracellular (left) and intercellular (right) metabolites significantly altered in PBM relative to PYG+ (positive Log2FC indicates higher levels in PBM). (A-B) *G. uro* was cultured anaerobically in EA-supplemented PBM or PYG+ for 48 h, followed by extraction and LC-MS analysis of intracellular and extracellular metabolites. **(C)** RNA-seq based pathway enrichment analysis of genes upregulated in PBM relative to PYG+. Bubble size represents the DEG count, and color intensity indicates the P value. **(D)** Volcano plot of RNA-seq data showing differentially expressed genes (DEGs) in *G. uro* grown in PBM relative to PYG+. Genes related to pyruvate metabolism and hydrogenase activity are highlighted in blue and orange, respectively. **(E)** Relative mRNA expression of pyruvate metabolism and hydrogenase genes. Data represent the fold change in gene expression in PBM relative to the PYG+ control, calculated using 2^-ΔΔCt^ method. *G. uro* was cultured anaerobically in EA-supplemented PBM or PYG+ for 48 h, followed by RNA isolation and reverse transcription. All data are presented as mean ± SEM (*n* = 3-5). DEGs were identified using DESeq2 with Benjamini-Hochberg FDR correction (FDR < 0.05); pathway enrichment was determined by hypergeometric test with FDR correction. Statistical significance for RT-qPCR data was assessed using Student’s t-test (*, P < 0.05; **, P < 0.01; ***, P < 0.001).

### Arginine-to-pyruvate shift induces metabolic and transcriptional reprogramming in *G. uro*

To elucidate the molecular mechanisms driving such outcomes, we performed integrated metabolomic and transcriptomic analyses. We first examined the metabolic shifts by comparing the intracellular and extracellular metabolomes of *G. uro* cultured in PBM versus PYG+ (both supplemented with EA). Principal coordinate analysis (PCA) revealed distinct clustering, indicating that PBM induces a fundamental physiological reconfiguration (**Fig. 3A**).

Untargeted metabolomics identified a broad range of significantly altered metabolites in both extracellular and intracellular fractions (**Fig. 3B**). Specifically, higher levels of arginine-related metabolites, including L-ornithine and L-citrulline, key intermediates of the arginine deiminase (ADI) pathway, as well as D-proline, which is metabolically linked to arginine catabolism, were detected in PYG+. The depletion of these metabolites in PBM confirms that arginine limitation effectively shuts down this energy-generating pathway.

To understand how *G. uro* compensates for this loss of arginine-based energy, we analyzed the transcriptional response using RNA-seq across four media conditions (PYG+, PYG+(+pyr), PYG+(-arg), PBM). Focusing on the PBM versus PYG+ comparison, we identified 1,371 differentially expressed genes (DEGs). Under arginine-limiting condition in PBM, the bacterium upregulated alternative energy-generating pathways. KEGG analysis revealed a robust enrichment of genes involved in pyruvate metabolism and TCA cycle (**Fig. 3C**). This transcriptional shift was visibly distinct in the volcano plot, which highlighted the specific upregulation of PFOR genes (*nifJ*, *porA*) alongside several hydrogenase genes, including the nickel-dependent hydrogenase, Ni/Fe hydrogenase, and maturation factors, such as *hypD* and the *HypC/HybG/HupF* family chaperone (**Fig. 3D and Supplementary Table 3**). Consistently, the PBM vs. PYG+(+pyr) comparison also showed an upregulation of pyruvate metabolism and hydrogenase genes in PBM (**Supplementary Fig. 4 and Supplementary Table 3)**, confirming that pyruvate utilization is suppressed when arginine is available.

RT-qPCR analysis confirmed this metabolic shift from arginine deamination to pyruvate oxidation and hydrogen production. We observed a significant induction of the pyruvate oxidizers *nifJ* (8.9 ± 2.0-fold) and *porA* (7.3 ± 0.6-fold) (**Fig. 3E**), which catalyze the conversion of pyruvate to acetyl-CoA. Meanwhile, essential hydrogenase determinants were strongly upregulated, including the maturation factor *hypB* (11.1 ± 3.0-fold) and structural subunits Hyd-L (3.8 ± 0.6-fold) and Hyd-S (2.5 ± 0.9-fold) (**Fig. 3E**). The oxidative decarboxylation of pyruvate generates excess reducing equivalents, primarily in the form of reduced ferredoxin (Fdred). To maintain redox balance, the upregulated membrane-bound hydrogenases facilitate the transfer of these electrons to protons (H^+^), resulting in the evolution of molecular hydrogen. Collectively, these findings demonstrate that PBM drives a coordinated metabolic reprogramming: the suppression of the ADI pathway necessitates the activation of the pyruvate-to-acetyl-CoA axis and subsequent molecular hydrogen evolution, which facilitates the reductive dehydroxylation required for urolithin production.

### Stable isotope tracer analysis validates pyruvate oxidation by *G. uro* in PBM

To validate the potentially upregulated pyruvate oxidation and its promoting role in hydrogen production as evidenced by transcriptomics data, we performed a ^13^C-isotopic tracing experiment using [3-^13^C]-pyruvate to characterize the metabolic fate of pyruvate. While the C-1 carbon of pyruvate is lost as CO2 during oxidative decarboxylation, the C-3 carbon is channeled into the acetyl-CoA pool via PFOR pathway. Specifically, utilizing C3-labelled tracer in this study enables us to directly monitor the incorporation of pyruvate-derived carbons into downstream metabolites, preventing carbon scrambling that may occur with a C-1 label.

Analysis of the extracellular medium revealed that the M+1 fraction of pyruvate was 82.0 ± 0.2% in the bacteria-free control and 72.9 ± 6.3% following a 48-h incubation with *G. uro* (**Fig. 4A**), indicating that a small fraction of unlabelled pyruvate, likely derived from other nutrient sources in the medium, was also present in the extracellular pool.

**Fig. 4.**
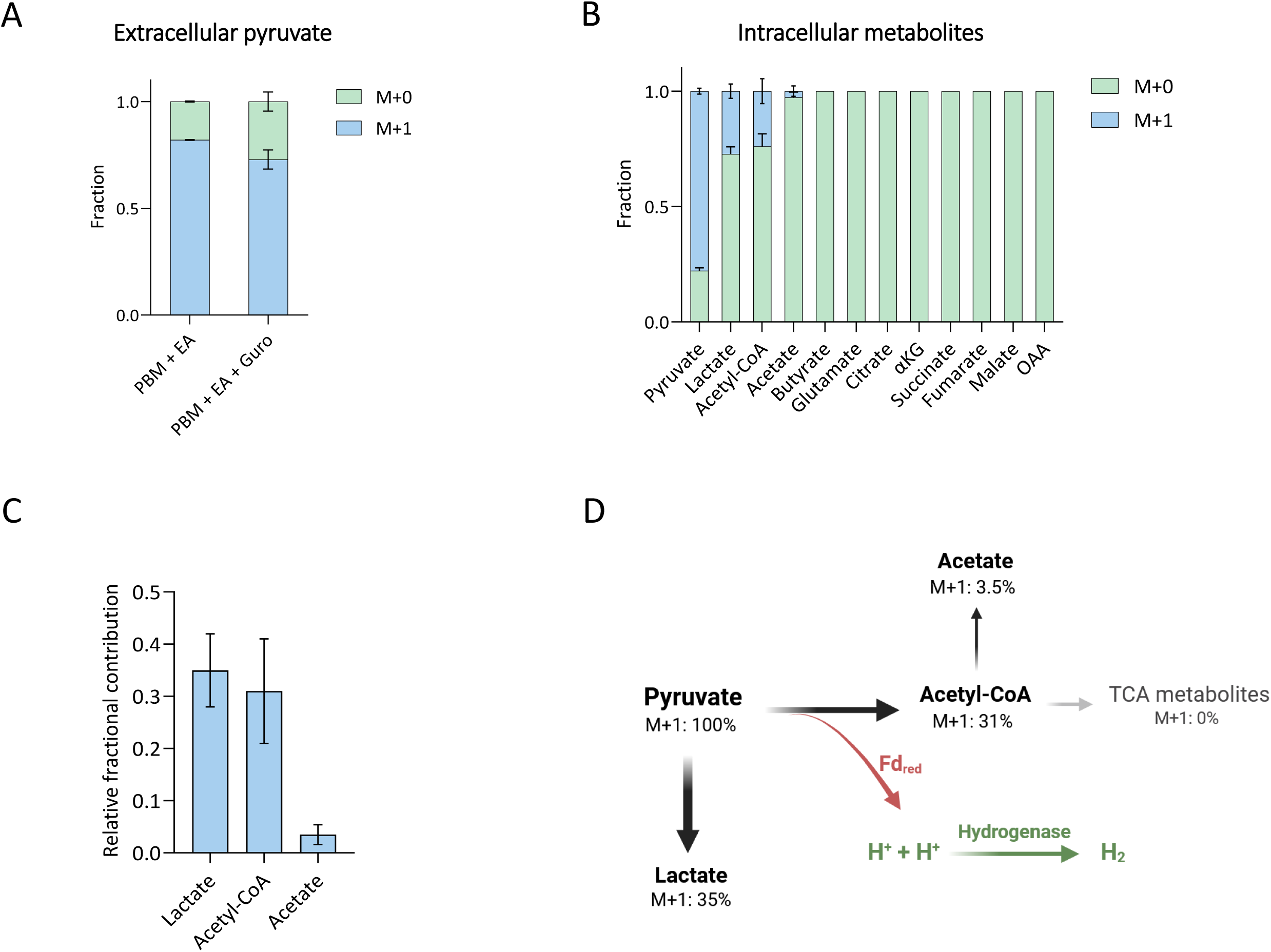
Stable isotope tracer analysis of pyruvate metabolism in *G. uro* using [3-^13^C]-pyruvate tracing combined with LC-MS. **(A)** Mass isotopologue distribution (M+0 and M+1 fractions) of extracellular pyruvate. **(B)** Mass isotopologue distributions of intracellular pyruvate, and pyruvate-derived organic acids. **(C)** Relative fractional contribution of [3-^13^C]-pyruvate to intracellular metabolite pools, calculated by normalizing the M+1 isotopologue fraction of each metabolite to the intracellular M+1 pyruvate fraction. **(D)** Schematic representation of metabolic partitioning and fate of pyruvate in *G. uro*. Pyruvate is predominantly distributed into lactate and acetyl-CoA. The conversion of pyruvate to acetyl-CoA yields reduced ferredoxin (Fdred), supplying electrons for hydrogen production. *G. uro* cultures in PBM were supplemented with EA and [3-^13^C]-pyruvate and incubated anaerobically for 48 h. All data are presented as mean ± SEM (*n* = 3). α-KG, α-ketoglutarate; OAA, oxaloacetate.

Intracellularly, M+1 pyruvate accounted for 77.9 ± 1.8% of the total pyruvate pool (**Fig. 4B**). In addition to pyruvate, isotopic labelling was directly detected in the intracellular pools of three downstream metabolites, with the M+1 fractions averaging 27.3 ± 5.4% for lactate, 24.0 ± 7.6% for acetyl-CoA, and 2.7 ± 1.2% for acetate. To accurately estimate the contribution of this labelled substrate to downstream metabolism, we calculated the relative fractional contribution by normalizing these values to the intracellular M+1 pyruvate pool. This analysis revealed that the [3-^13^C]-pyruvate tracer contributed to 35.0 ± 5.0% of the intracellular lactate pool and 31.0 ± 11.0% the acetyl-CoA pool, while contributing minimally (3.5 ± 1.8%) to the intracellular acetate pool. The substantial fractional contribution observed in the acetyl-CoA pool directly validates the upregulated PFOR-mediated tracing. Extracellular analysis further detailed the metabolic fate of the pyruvate, where the M+1 isotopic fractions for lactate and acetate were measured at 31.3 ± 5.2% and 1.9 ± 1.1%, respectively (**Supplementary Fig. 5**).

Despite the efficient incorporation of the tracer into acetyl-CoA, there was no detectable flux into the TCA cycle (**Fig. 4B**). The absence of labeling in glutamate and butyrate suggests that pyruvate-derived acetyl-CoA is not further metabolized through these oxidative or biosynthetic routes. Crucially, the substantial contribution of pyruvate to the acetyl-CoA pool via PFOR generates Fdred, providing the essential reducing power for hydrogen evolution through hydrogenase activity (**Fig. 4D**).

### Hydrogen promotes multi-step EA-to-urolithin conversions

Inspired by a recent study reporting that hydrogen promotes the 21-dehydroxylation of biliary corticoids into progestins by *E. lenta* ^22^, we hypothesized that hydrogen is a key driver of the enhanced dehydroxylation activity in *G. uro*. Headspace GC-TCD analysis confirmed that *G. uro* actively produces hydrogen gas in PBM (∼3.01 µmol), whereas hydrogen production was significantly lower in PYG+ (∼0.07 µmol; P < 0.001) after 72-h incubation (**Fig. 5A**). Consistent with this, a similar trend was observed when quantifying dissolved hydrogen directly in the culture media using a methylene blue-based titration method, further validating the enhanced hydrogen production in PBM (**Supplementary Fig. 6**). Our transcriptomic data also indicates a marked upregulation of hydrogenase genes in PBM (Fig. 3).

**Fig. 5.**
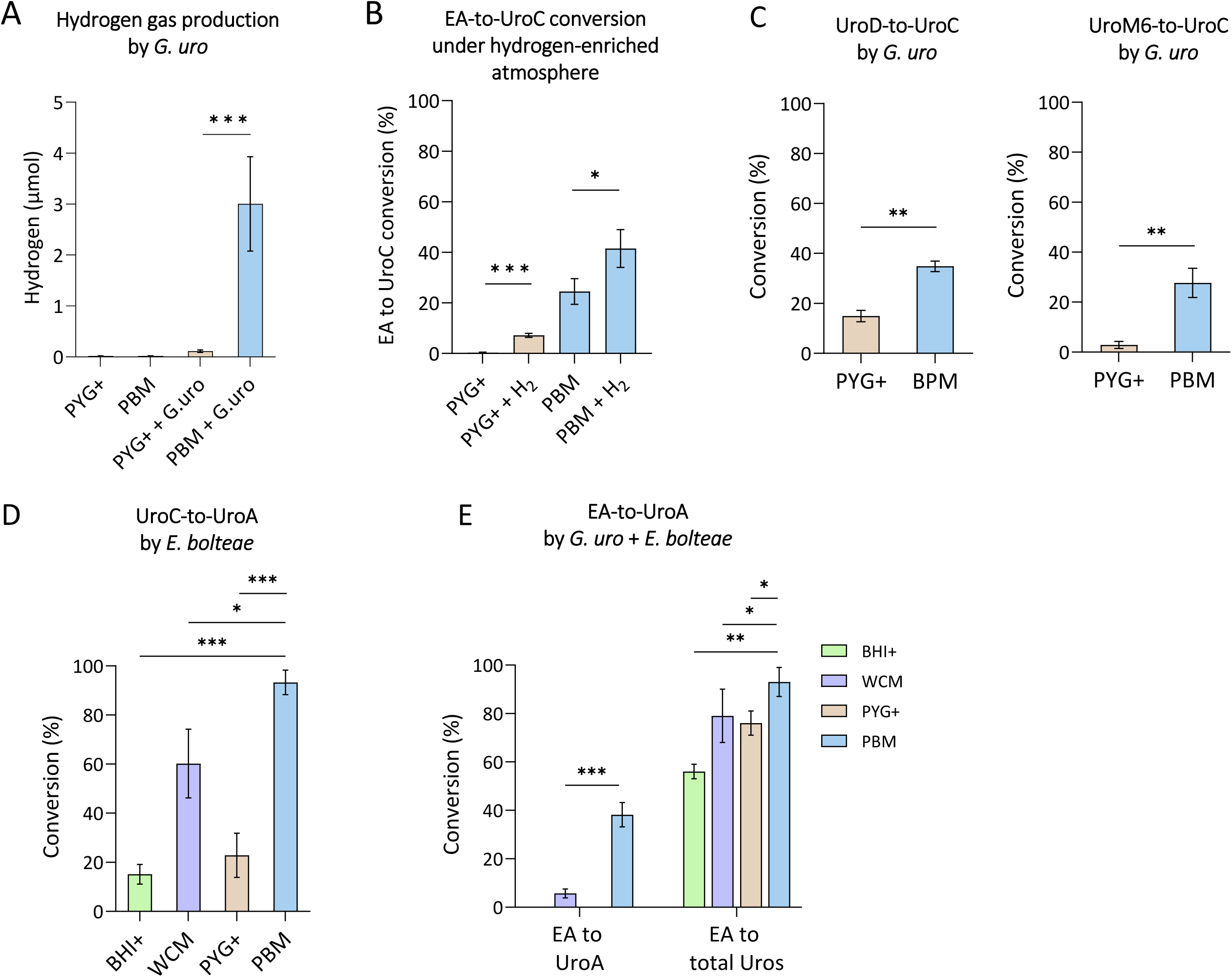
Hydrogen enhances gut bacterial dehydroxylation of urolithins. **(A)** Hydrogen gas production by *G. uro* cultured in PYG+ and PBM after 3 days of fermentation in airtight serum bottles. Hydrogen was measured in the bottle headspace using GC-TCD. **(B)** EA-to-UroC conversion by *G. uro* under a hydrogen-enriched atmosphere established using hydrogen tablets compared to standard anaerobic atmosphere in PYG+ and PBM. **(C)** Conversion of UroD to UroC and UroM6 to UroC by *G. uro*. **(D)** Conversion of UroC to UroA by *E. bolteae.* **(E)** Conversion of EA to UroA by *G. uro* and *E. bolteae* co-culture. For all bioconversion assays, culture media were supplemented with a phenolic substrate (40 µM) and then incubated anaerobically with bacteria for 72 h. Data are presented as mean ± SEM (*n* = 3). Statistical significance was assessed using one-way ANOVA followed by Tukey’s post hoc test (multiple groups) or Student’s t-test (two groups) (*, P < 0.05; **, P < 0.01; ***, P < 0.001).

To determine if hydrogen availability is the bottleneck in nutrient media, we supplemented PYG+ with exogenous hydrogen using hydrogen-producing tablets. Under a hydrogen-enriched atmosphere, EA-to-UroC conversion in PYG+ increased significantly from 0.5% to 7.2%. Furthermore, adding excess hydrogen to PBM further boosted conversion yields to 41.5%, confirming that dehydroxylation is hydrogen-dependent (**Fig. 5B**). Moreover, a higher level of hydrogen enhanced UroC production in both media (P < 0.05) (**Supplementary Fig. 7**).

Having established that PBM naturally provides the reducing conditions necessary to drive these upstream dehydroxylations, we confirmed that this hydrogen-driven enhancement extended beyond the initial EA-to-UroC pathway, significantly improving the conversion of key intermediates such as UroD and UroM6 into UroC (**Fig. 5C**). We next investigated whether this optimized environment could also enhance the final downstream step catalyzed by *E. bolteae*. When supplied directly with UroC as a substrate, *E. bolteae* alone achieved near-complete conversion to UroA in PBM (93.4 ± 5%), significantly outperforming standard PYG+ media (**Fig. 5D**). Finally, to determine if PBM could support the complete conversion of EA to the final end-product, *G. uro* and *E. bolteae* were co-cultured. In this consortium, PBM enabled the robust, continuous production of UroA directly from EA (38.3 ± 5%), whereas conversion was negligible in controls (**Fig. 5E**). These results demonstrate that PBM potentially creates a hydrogen-rich environment that unlocks the full catalytic potential of the urolithin-producing consortium.

### Co-culturing with hydrogen-producing *Bacteroides* enhances urolithin production

To confirm that cross-feeding hydrogen can drive urolithin dehydroxylation by *G. uro*, we co-cultured it with three *Bacteroides* species exhibiting distinct hydrogen-producing profiles. GC-TCD analysis confirmed that *B. vulgatus* is a high hydrogen producer and *B. thetaiotaomicron* is a moderate producer, whereas *B. stercoris* produces negligible hydrogen (**Fig. 6A**).

**Fig. 6.**
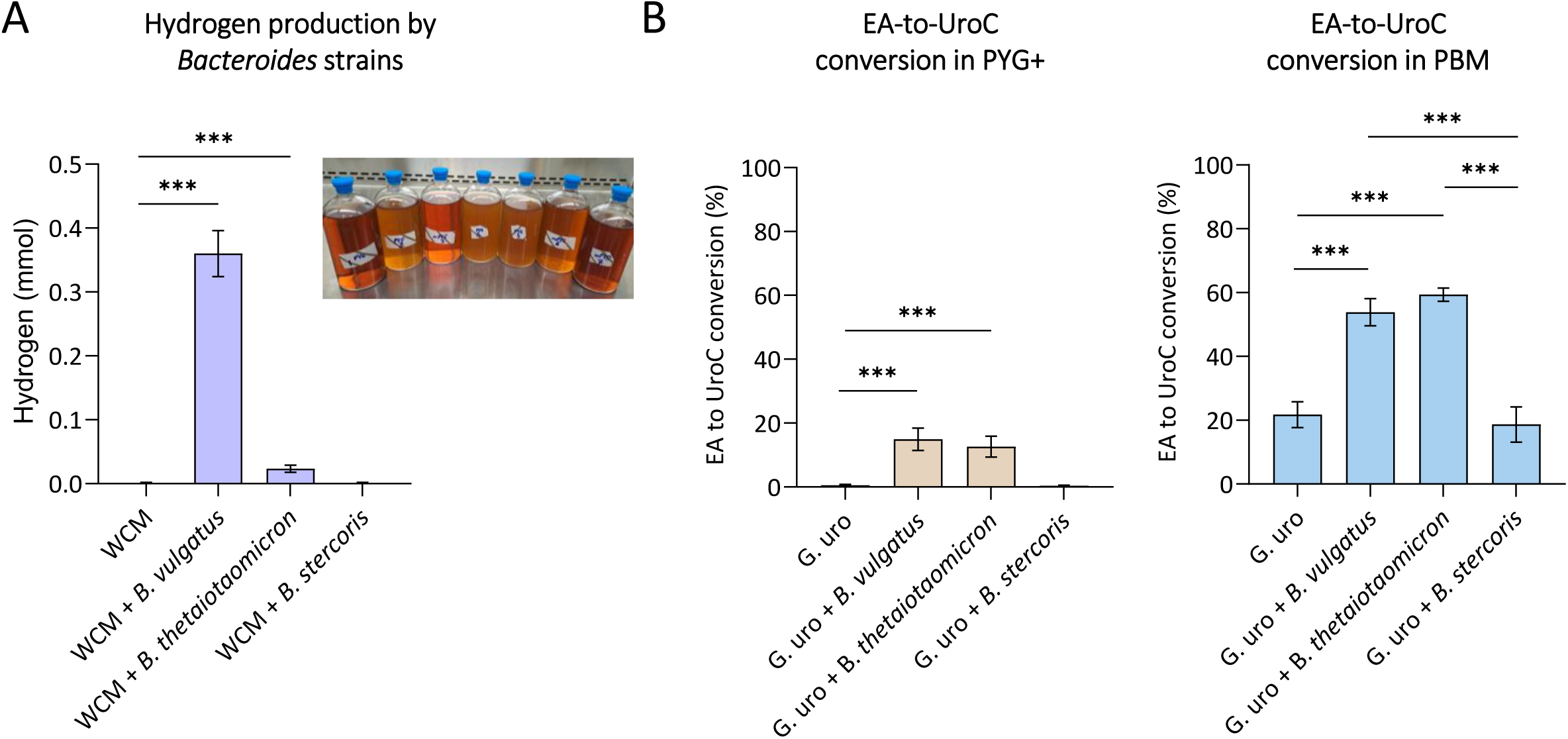
Co-culturing *G. uro* with hydrogen-producing bacteria enhances EA-to-urolithin conversion. **(A)** Hydrogen production by individual *Bacteroides* strains (*B. thetaiotaomicron* ATCC 29148, *B. vulgatus* ATCC 8482, and *B. stercoris* DA816) cultured in WCM medium. **(B)** EA-to-UroC conversion by *G. uro* co-cultured with each *Bacteroides* strain in PYG+ or PBM supplemented with EA (40 µM). *G. uro* was co-cultured with the indicated strains under anaerobic conditions for 48 h. Data are presented as mean ± SEM (*n* = 3). Statistical significance was assessed using one-way ANOVA followed by Tukey’s post hoc test (*, P < 0.05; **, P < 0.01; ***, P < 0.001).

Co-culturing *G. uro* with *B. vulgatus* or *B. thetaiotaomicron* significantly increased EA-to-UroC conversion (P < 0.05), whereas the non-hydrogen producer *B. stercoris* failed to promote any significant conversion in PYG+ (**Fig. 6B**). Furthermore, even in the optimal PBM environment, the presence of hydrogen-producing partners generated a synergistic effect, doubling the conversion yield compared to *G. uro* monocultures (P < 0.05) (**Fig. 6B**). Ultimately, these findings demonstrate that *G. uro* can utilize exogenous hydrogen supplied by microflora members to activate the EA metabolic cascade and drive robust urolithin production.

### Pyruvate supplementation with arginine restriction enhances EA-to-urolithin conversion in mice

To evaluate the physiological relevance of our *in vitro* findings, we investigated whether increasing the dietary pyruvate intake could promote EA-to-urolithin bioconversion in wild-type C57BL/6J mice naturally devoid of *G. uro* ^16^. Mice were orally dosed with live *G. uro* and maintained on a high-pyruvate (arginine-limited) or high-arginine dietary regimen, designed to mimic the nutritional conditions of PBM and PYG+, respectively. Then EA was administered to mice before urine and feces were collected over 48 hours for EA and urolithin analysis.

The mean total urinary excretion of UroC was significantly higher in mice receiving the pyruvate-supplemented diet (0.89 ± 0.15 nmol) compared to the high-arginine group (0.33 ± 0.10 nmol, P < 0.01) (**Fig. 7C**). Furthermore, mice in the PBM group exhibited slightly higher excretion of UroA (0.96 ± 0.09 nmol, P > 0.05). These analyses indicate that dietary pyruvate effectively supports the initial dehydroxylation steps in the host environment.

**Fig. 7.**
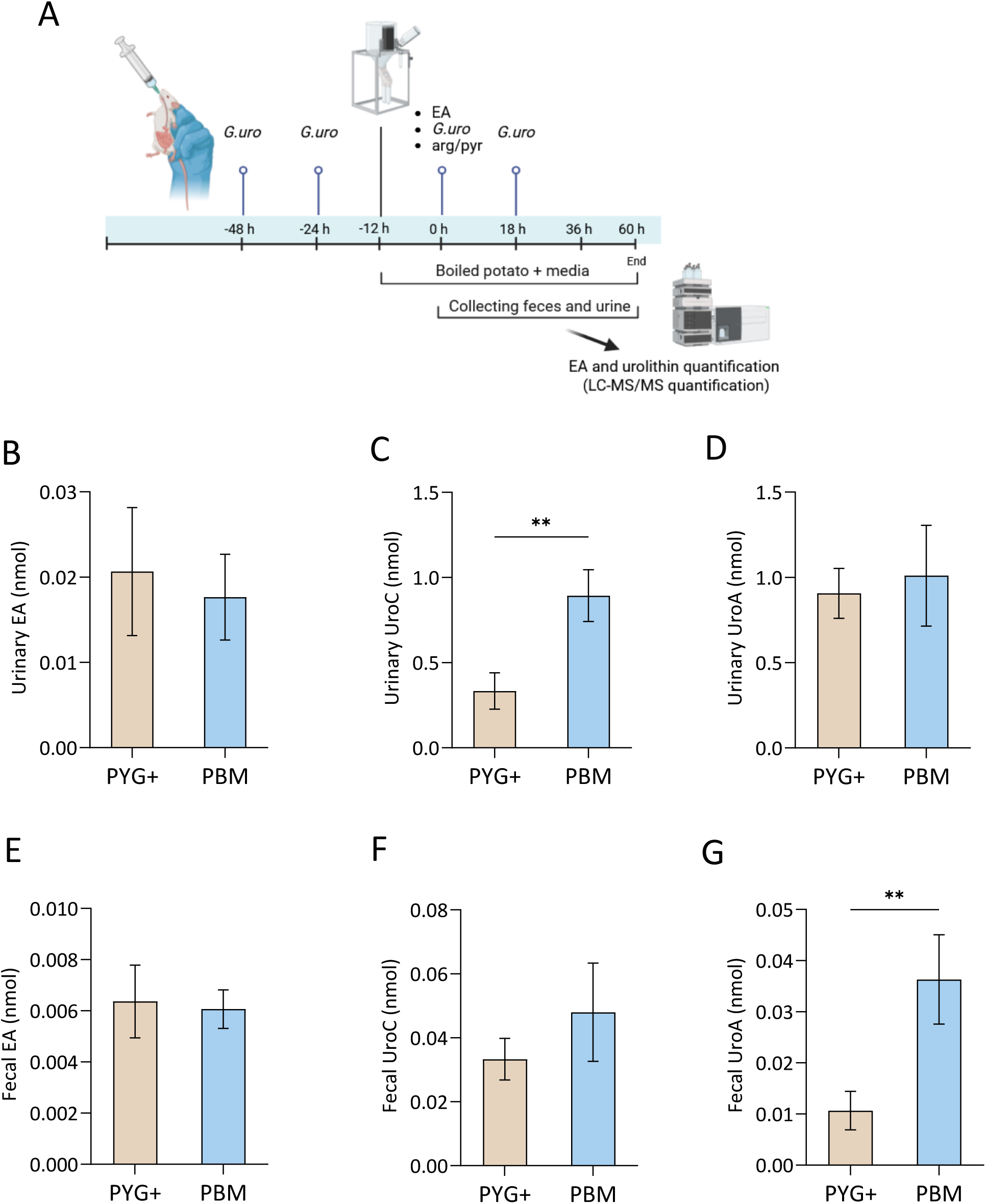
High dietary pyruvate with arginine limitation enhances EA-to-urolithin conversion in mice. **(A)** Animal experimental design. **(B–D)** Total excretion of EA and urolithins in urine over 48 hours post-gavage of EA. **(E–G)** Total excretion in feces over 48 hours post-gavage. Mice were administered *G. uro*, EA, and maintained on a high-arginine PYG+ regimen or a high-pyruvate PBM regimen. All data are presented as mean ± SEM (*n* = 6). Statistical significance was assessed using Student’s t-test (*, P < 0.05; **, P < 0.01).

Consistent with the urinary data, fecal analysis demonstrated a significant increase in the production of the final end-product. Mice on the high-pyruvate regimen excreted significantly higher levels of UroA (0.036 ± 0.005 nmol) compared to the control group (0.011 ± 0.002 nmol) (P < 0.05) (**Fig. 7G**). A similar upward trend was observed for fecal UroC in the PBM group (0.048 ± 0.005 nmol) versus the PYG+ (0.033 ± 0.002 nmol); however, this increase was not statistically significant (P > 0.05). Collectively, these data demonstrate that dietary pyruvate reprograms gut microbial metabolism *in vivo*, promoting the conversion of EA into bioavailable urolithins.

### PBM enhances bacterial conversion of polyphenols with pyrogallol and catechol structural units

Given the remarkable EA metabolizing effects conferred by PBM, we hypothesized that similar catabolic-promoting activities could be reproduced in other gut bacteria. Since *E. lenta* possesses similar decarboxylation and dehydroxylation capabilities for polyphenols containing pyrogallol and catechol structural units ^14^, we further tested the impact of PBM on these broader biotransformations.

First, we investigated the conversion of epicatechin by *E. lenta*. PBM significantly enhanced the conversion efficiency to 66.4 ± 2.9%, significantly higher than the rates observed in BHI+, WCM, and PYG+ (**Fig. 8A**). Next, we assessed the metabolism of caffeic acid by *G. uro*. Consistent with our previous findings, the pyruvate-enriched environment approximately doubled the conversion rate compared to that in PYG+ (**Fig. 8B**). These results suggest that PBM creates a favorable environment that promotes diverse biotransformations in multiple polyphenol-degrading gut bacteria.

**Fig. 8.**
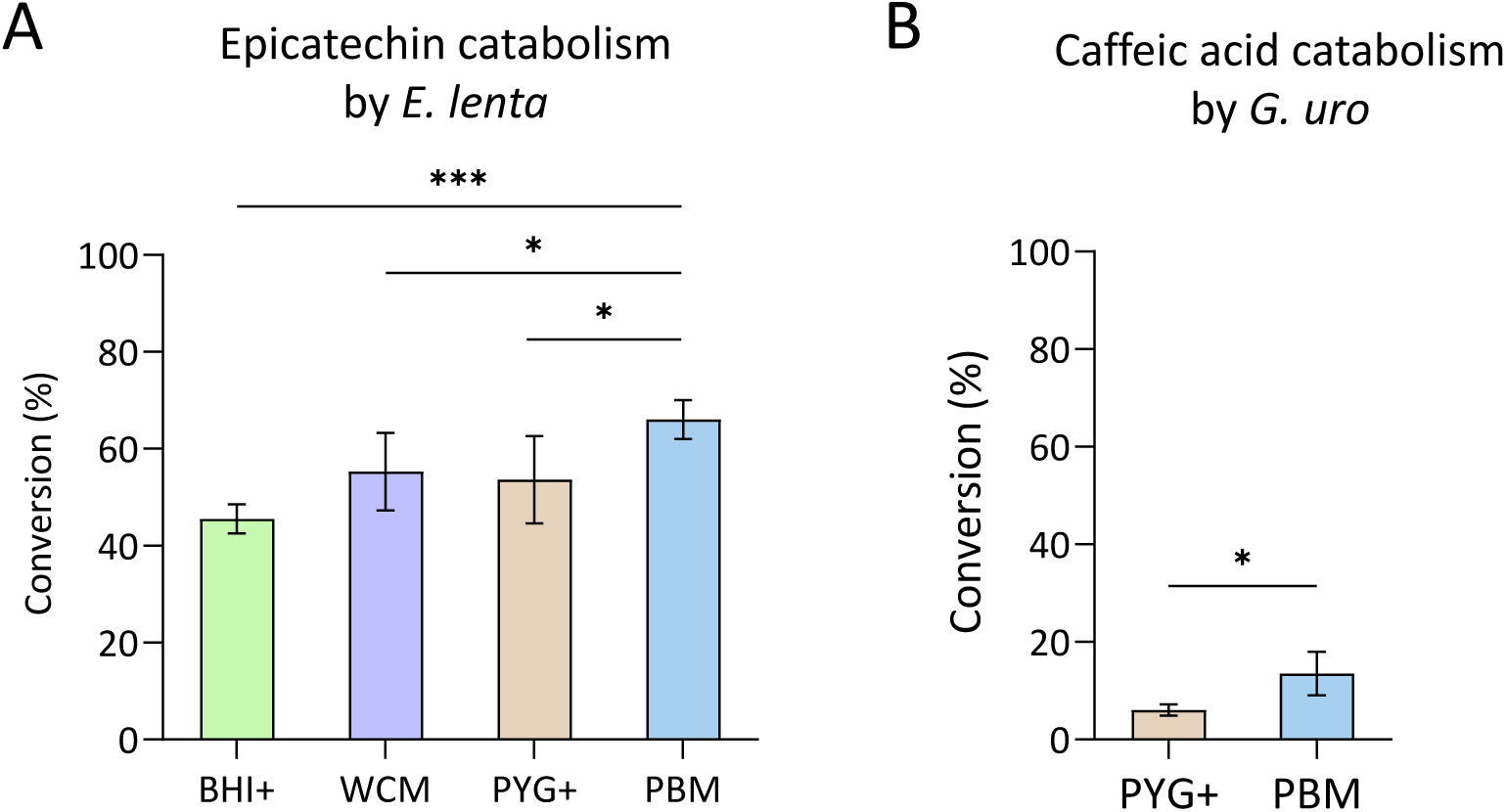
PBM enhances polyphenol bioconversion by multiple gut bacterial species. **(A)** Hydrolysis of epicatechin by *E. lenta* in BHI+, WCM, PYG+, and PBM. **(B)** Hydrolysis of caffeic acid by *G. uro* in PYG+ and PBM. Data are presented as mean ± SEM (*n* = 3). Statistical significance was assessed using one-way ANOVA followed by Tukey’s post hoc test (multiple groups) or Student’s t-test (two groups) (*, P < 0.05; **, P < 0.01; ***, P < 0.001).

## DISCUSSION

Recently, significant progress has been made in understanding the enzymatic repertoire involved in EA-to-urolithin metabolism ^15, 23^, but the environmental factors influencing this bioconversion remained underexplored. Herein, we report that shifting from an arginine-rich to a pyruvate-rich environment substantially enhances EA-to-urolithin conversion by gut bacteria, with hydrogen gas serving as a key driver for dehydroxylation. Application of the newly developed medium, PBM, which enhanced the catabolism of various polyphenols containing pyrogallol and catechol moieties, reveals that pyruvate-driven hydrogen production is the pivotal mechanism mediating reductive dehydroxylation in polyphenol-degrading gut bacteria.

Previously, we observed that while EA and *G. uro* co-administration to mice resulted in remarkable UroC production ^16^.However, such efficient EA bioconversion was not observed when *G. uro* was cultured in standard growth media *in vitro*, despite its active proliferation. To identify the factors limiting *in vitro* EA bioconversion, we examined the nutrient compositions of the commonly used nutrient media for culturing strict anaerobic gut bacteria. The presence of glucose is common in most standard growth media including PYG+ and BHI+, but *G. uro* is unable to metabolize it ^21^, and its removal does not influence its growth nor metabolic efficiency (**Supplementary Fig. 8**). Instead, arginine fuels central carbon metabolism in *G. uro* (**Fig. 2C**) and *E. lenta* ^20^, predominantly through the ADI pathway, promoting fast biomass accumulation (**Fig. 3C**). However, such rapid growth in arginine-rich conditions did not favor EA bioconversion (**Fig. 1C**). Another standard medium that supports the growth of *Eggerthellaceae* species is WCM, which contains no glucose and low arginine content, but provides pyruvate as the major carbon source, contrasting the nutrient profiles in other standard media tested. Interestingly, although *G. uro* proliferation was delayed as pyruvate appears to be a less efficient energy substrate than arginine, the average EA-to-urolithin conversion rate was faster than those in PYG+ and BHI+ (**Supplementary Fig. 9**). We thus hypothesize that with limited arginine availability, *G. uro* undergoes a metabolic shift that activates alternative pathways, such as carbon fixation pathways to generate acetate from CO₂ (**Fig. 3C**), allowing other reductive processes (e.g., EA/urolithin dehydroxylation) to take place in the meantime ^24^. Of note, the carbon fixation pathways have also been identified as the major energy-conservation strategy in *E. lenta*, which belongs to the same family, *Eggerthellaceae*, as *G. uro* ^20, 25^.

As demonstrated by transcriptomics and RT-qPCR analyses, *G. uro* metabolized pyruvate through the PFOR pathway in PBM (**Fig. 3D-F**), with concurrent upregulation of hydrogenase genes, including both maturation factors and structural subunits (**Fig. 3D and 3F**). Although these hydrogenase genes have not yet been well characterized, their sequence homology suggests that they belong to the [NiFe]-hydrogenase family ^26^. Consistently, stable isotope tracer analysis using the [3-^13^C]-pyruvate confirmed PFOR activity by direct detection of labelled acetyl-CoA (**Fig. 4B**). This reaction generates reduced ferredoxin, which serves as the electron donor for hydrogenases to drive hydrogen evolution ^27^. The functional activation of hydrogenases was further supported by the detection of substantial hydrogen gas in the headspace of PBM cultures, but not in PYG+ (**Fig. 5A**). Consistently, a recent work by McCurry et al. reported that hydrogen gas promotes corticosteroid 21-dehydroxylation in *E. lenta* ^22^. In addition to the roles of endogenous hydrogen, we also demonstrated that exogenous hydrogen supplementation through using hydrogen-releasing tablets or co-culturing with hydrogen-producing *Bacteroides* spp. readily alleviated the repressive conditions in PYG+ and further enhanced UroC production in PBM (**Fig. 4B**). Within the gut microflora, this hydrogen-rich environment can be generated through microbial cross-feeding, wherein fiber-degrading bacteria, such as *Bacteroides* and *Clostridium* species, produce substantial hydrogen gas as a fermentative byproduct ^28, 29^. These findings underscore the essential roles of hydrogen gas in *Gordonibacter*-mediated EA-to-urolithin conversion.

Our *in vitro* findings reveal a clear dissociation between EA-metabolizing bacterial biomass and metabolic capacity, suggesting that the conventional probiotic strategy of administering high doses of urolithin-producing bacteria may not be sufficient if the host gut environment channels them toward arginine catabolism rather than polyphenol transformation. Moving from *in vitro* to *in vivo* settings, our proof-of-concept mouse study corroborates this nutritional cue-driven mechanism: dietary pyruvate supplementation along with arginine limitation significantly increased UroC and UroA production compared to the arginine-rich, pyruvate-deficient controls (**Fig. 7)**. This mechanism is further supported by a recent study from Sinha et al., which demonstrated that gut microbial tryptophan metabolism is similarly governed by substrate-dependent regulation, driven by dietary fiber and cross-feeding, rather than the abundance of the responding bacteria ^30^.

The observations that PBM enhances not only EA-to-urolithin conversion but also epicatechin catabolism by *E. lenta* and caffeic acid conversion by *G. pam* (**Fig. 8**) suggest that hydrogen-driven dehydroxylation may represent a shared mechanism across diverse polyphenol classes, operating within and supported by bacterial cross-feeding networks. This aligns with the emerging view that catechol dehydroxylases, either the DMSO reductase family (characterized in *Gordonibacter* and *Eggerthella*) or the xanthine oxidase family (characterized in *Enterocloster*), universally require reduced electron carriers ^14, 15, 31^. Consistent with this strict dependence on available reducing power, introducing a competing electron sink via supplementing additional hemin significantly impaired conversion (**Supplementary Fig. 10**). Ultimately, although catechol dehydroxylases exhibit distinct substrate specificities and regioselectivities ^14, 15^, their functional activity depends on a sustained supply of reducing equivalents under anaerobic conditions.

While this study identifies hydrogen as a primary regulator of polyphenol dehydroxylation, we acknowledge several limitations which require further investigations. First, the precise biochemical mechanism by which hydrogen promotes dehydroxylase activity, whether through direct electron donation to the molybdenum cofactor or indirect regeneration of reduced ferredoxin, remains to be elucidated. Second, although interspecies hydrogen transfer occurs readily *in vitro*, the broader ecological competition governing endogenous versus community-supplied hydrogen utilization *in vivo* represents a critical frontier that we are currently exploring. Third, the durability and host-level consequences of sustained dietary pyruvate supplementation require further evaluation in chronic studies.

In this study, we demonstrate that efficient EA-to-urolithin conversion by *G. uro* requires a nutritional shift from arginine-driven growth to pyruvate-driven hydrogen production, achieved through our developed PBM. This hydrogen serves as the critical engine driving the urolithin dehydroxylation cascade, enabling robust UroC production and, in co-culture with *E. bolteae*, UroA formation. Supported by our *in vivo* mouse model and evidence of exogenous hydrogen cross-feeding, this dehydroxylation emerges as a cooperative community trait. Our findings establish hydrogen as a key driver of microbial polyphenol metabolism. This discovery suggests that modulating the dietary arginine-to-pyruvate ratio can effectively enhance gut microbiota’s capacity to produce bioactive metabolites. Given the well-documented anti-inflammatory, anti-obesity, and anti-aging effects of UroA, this targeted dietary strategy positions these metabolites as highly valuable pharmaceutical and nutraceutical targets.

## METHODS

### Chemicals and reagents

All chemicals and reagents used in this study were of analytical or HPLC grade. Ellagic acid (EA) was purchased from Macklin Biochemical Co., Ltd. (Shanghai, China). Urolithin C (UroC), urolithin M6 (UroM6), urolithin D (UroD), and urolithin M7 (UroM7) were obtained from Molcore (Nanjing, China). 6,7-Dihydroxycoumarin was purchased from Yuanye Bio-Technology (Shanghai, China). Caffeic acid and epicatechin were obtained from Aladdin Biochemical Technology Co., Ltd. (Shanghai, China). Short-chain fatty acid (SCFA) standards were purchased from Macklin Biochemical Co., Ltd. (Shanghai, China). Solvents for LC-MS analyses included methanol (Duksan, Ansan, South Korea), formic acid (Macklin, Shanghai, China), and ethyl acetate (Anaqua Chemicals Supply, Wilmington, OH, USA). Stable isotope-labelled compounds included sodium [3-^13^C]-pyruvate (sodium 2-oxopropanoate-¹³C) from MedChemExpress (Monmouth Junction, NJ, USA) and succinate-d₄ from Cambridge Isotope Laboratories (Tewksbury, MA, USA).

Derivatization reagents included 1-ethyl-3-(3-dimethylaminopropyl)carbodiimide (EDC) from Macklin Biochemical Co., Ltd. (Shanghai, China), O-benzylhydroxylamine (OBHA) from Yuanye Bio-Technology (Shanghai, China), and pyridine from Aladdin Biochemical Technology Co., Ltd. (Shanghai, China). Media components and minerals included dipotassium phosphate (Aladdin, Shanghai, China), tween 80 (Aladdin, Shanghai, China), L-arginine (Biopik, North Brunswick, NJ, USA), resazurin (Macklin, Shanghai, China), lysozyme (Aladdin, Shanghai, China), protease inhibitors (EDTA-free; Roche Diagnostics, Mannheim, Germany), vitamin K and hemin (Thermo Scientific, Waltham, MA, USA), and L-cysteine (Aladdin, Shanghai, China). Dissolved hydrogen concentrations were measured using a methylene blue-platinum titration reagent (H_2_Blue; Kueysing, China). For LC-MS analyses, stock solutions of EA and urolithins were initially prepared in pure methanol. Working standard solutions were subsequently prepared by diluting the stock solutions with 80% methanol containing 0.1% (v/v) formic acid, unless otherwise indicated.

### Bacterial strains and culture conditions

The bacterial strains used in this study included *Gordonibacter urolithinfaciens* DSM 27213 (*G. uro*), *G. pamelaeae* DSM 19378 (*G. pam*), *Eggerthella lenta* DSM 2243 (*E. lenta*), and *Enterocloster bolteae* DSM 15670 (*E. bolteae*), all obtained from the Leibniz Institute DSMZ-German Collection of Microorganisms and Cell Cultures (Braunschweig, Germany). In addition, three *Bacteroides* strains, *B. thetaiotaomicron* ATCC 29148, *B. vulgatus* ATCC 8482, and *B. stercoris* DA816, were purchased from the American Type Culture Collection (ATCC, Manassas, VA, USA). The identity of all bacterial strains was validated by 16S rRNA gene sequencing prior to use.

Four distinct standard growth media were used in this study. (1) WCM: Wilkins-Chalgren Medium (Neogen, Lansing, MI, USA); (2) BHI+: Brain heart infusion (BHI; Hopebio, Shandong, China) supplemented with 5% (w/v) L-arginine monohydrochloride and 0.05% (w/v) L-cysteine monohydrochloride; (3) PYG+: peptone yeast glucose (PYG; Hopebio, Shandong, China) supplemented with 2 g/L dipotassium phosphate, 1 mL/L tween 80, 5 g/L L-arginine, 1 mg/L resazurin, 1 mg/L vitamin K, 5 mg/L hemin; (4) PBM, PhenolBoost Medium – A novel formulation developed in this study, consisting of PYG+ ingredients with L-arginine replaced by 1 g/L sodium pyruvate.

Frozen bacterial stocks were activated by streaking onto Columbia sheep blood agar plates (Hopebio, Shandong, China) and incubating anaerobically for three days at 37 °C, followed by transferring single colonies into WCM broth. Most bacterial procedures were performed inside an anaerobic workstation (Concept 400, Ruskinn Technology Ltd., Bridgend, UK) under an atmosphere of 5% hydrogen, 10% carbon dioxide, and nitrogen as the balance. Anaerobic jars equipped with anaerobic gas-generating sachets (Thermo Scientific, Waltham, MA, USA) were used for bacterial incubation.

### Evaluating conversion of EA, urolithins and other phenolic compounds

#### EA-to-urolithin conversion by *Gordonibacter*

The ability of *Gordonibacter* spp. to convert EA to urolithins was assessed in BHI+, WCM, and PYG+. Briefly, a 72-hour culture of *G. uro* or *G. pam* in WCM was diluted 1:100 into 250 µL of the respective bacterial media in 96-well plates, followed by EA supplementation to a final concentration of 40 µM. Control cultures received an equivalent volume of dimethyl sulfoxide (DMSO) instead of EA to establish baseline measurements for conversion calculations. The bacterial cultures were then incubated anaerobically at 37 °C for 72 h. Subsequently, the fermented cultures were collected for EA and urolithin extraction. In addition, the effect of EA concentrations (20, 40, 60 µM) and *G. uro* inoculum size (6.5, 7.5, 8.5 log CFU/mL) were examined in PYG+ medium as described above. All experiments were conducted in triplicate.

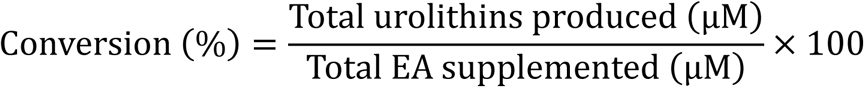

#### Conversion of urolithin intermediates

The biotransformation of urolithin intermediates was tested using multiple strains and substrates. *G. uro* was cultured in PBM and PYG+ with 40 µM UroD or UroM6 to assess conversion to UroC. *G. pam* was tested for EA-to-UroC conversion. *E. bolteae* was examined with 40 µM UroC to evaluate its conversion to UroA, while *G. uro* and *E. bolteae* co-cultures (1:1 ratio) were supplemented with 40 µM EA to assess UroA production capacity in PBM versus PYG+. All cultures were incubated anaerobically at 37°C for 72 h prior to extraction.

### EA-to-urolithin conversion by *G. uro* lysate

A 72-hour culture of *G. uro* in WCM was diluted 1:100 into 10 mL of fresh PYG+ or PBM and supplemented with 40 µM EA to induce enzyme expression. Cultures were incubated anaerobically for 72 h at 37°C. Bacterial cells were harvested by centrifugation (8,500 × *g*, 5 min) and washed with pre-reduced PBS. The cell suspensions were normalized to total protein content determined by the bicinchoninic acid (BCA) assay (Thermo Scientific, Waltham, MA, USA). Harvested cells were resuspended with 1 mL of pre-reduced lysis buffer (20 mM Tris-HCl, 500 mM NaCl, 10 mM MgSO₄, 10 mM CaCl₂, pH 7.5) containing protease inhibitors (EDTA-free). Lysozyme was added to a final concentration of 5 mg/mL, and the suspension was incubated for 60 min at 37°C. The mixture was centrifuged at 17,000 × *g* for 4 min to pellet insoluble debris. A 0.9 mL aliquot of the clarified lysate was transferred to a fresh tube, maintained on ice, and supplemented with 40 µM EA. The reaction mixture was incubated anaerobically for 48 h at 37°C before extraction. All steps, excluding centrifugation, were performed under strict anaerobic conditions.

#### Metabolism of other phenolic compounds

The conversion of caffeic acid (100 µM) *by G. uro* was compared between PYG+ and PBM, and the metabolism of epicatechin (100 µM) by *E. lenta* was assessed in BHI+, WCM, PYG+, and PBM. All fermentation experiments were conducted according to the procedures detailed above.

#### Extraction of urolithins and phenolic compounds

Extraction was performed as previously described ^16^. Briefly, 250 µL fermented samples were extracted with ethyl acetate containing 1.5% (v/v) formic acid. 6,7-dihydroxycoumarin was employed as an internal standard (IS) at a concentration of 0.1 µg/mL. The extraction was repeated three times, the top organic layers were combined and dried using a SpeedVac concentrator (Thermo Scientific SPD410DDA, USA). The residue was reconstituted with 80% methanol containing 0.1% formic acid. The precipitates were removed by centrifugation at 17,000 *g* for 10 min prior to LC–MS/MS analysis.

#### LC-MS/MS quantification of phenolic metabolites

LC-MS/MS analysis was performed using an ExionLC™ AD series UHPLC system coupled to a QTRAP® 6500+ mass spectrometer (Sciex, Framingham, MA, USA) equipped with an electrospray ionization (ESI) source. Chromatographic separation was achieved using an ACQUITY UPLC HSS T3 column (100 × 2.1 mm, 1.8 µm; Waters, Milford, MA, USA) maintained at 40 °C. The mobile phases consisted of water containing 0.1% formic acid (A) and acetonitrile containing 0.1% formic acid (B). Detailed chromatographic and MS parameters are provided in **Supplementary Methods**.

### Evaluating the roles of arginine and pyruvate on EA-to-urolithin conversion

#### Effects of pyruvate and arginine concentrations on conversion

To test the effect of arginine and pyruvate on EA metabolism, four specific media were used: (1) PYG+; (2) PYG+ without arginine (PYG+[-arg]); (3) PYG+ supplemented with 1 g/L pyruvate (PYG+[+pyr]); (4) PYG+ without arginine but supplemented with pyruvate (PBM). These media were inoculated with *G. uro* and supplemented with 40 µM EA as described above. Additionally, to optimize component levels, the effects of varying concentrations of pyruvate (0.5, 1.0, and 1.5 g/L) and arginine (2.5, 5.0, and 7.5 g/L) were assessed in PBM and PYG+, respectively.

To analyze the reaction kinetics, PBM cultures were inoculated with *G. uro* and supplemented with EA under the same conditions described above. Aliquots were collected at 0, 24, 48, and 72 h to monitor EA hydrolysis and urolithin production. In all experiments, metabolites were extracted and analyzed via LC-MS as described above.

#### Effects of pyruvate and arginine concentrations on bacterial growth

To assess the bacterial growth kinetics, freshly cultured *G. uro* in WCM was diluted 1:100 into 5 mL of PYG+, PYG+(-arg), PYG+(+pyr), and PBM, each supplemented with 40 µM EA. Bacterial enumeration was performed at 0, 24, 48, 72, and 96 h by plating serially diluted aliquots onto Columbia sheep blood agar plates. Colonies were counted after anaerobic incubation to determine CFU/mL.

### Assessing pyruvate and arginine consumption

#### Sample preparation, extraction, and derivatization

To determine the consumption rates of arginine and pyruvate nutrients, samples were collected from the *G. uro* growth experiments described above. For arginine and pyruvate extraction, 500 µL of fermented culture was centrifuged (8,500 × *g*, 5 min) to remove cells. The supernatants were mixed with 600 µL of 80% methanol containing succinate-d4 (0.1 µg/mL) as an IS. The mixtures were centrifuged at 17,000 × *g* for 10 min, and the collected supernatants were dried using a SpeedVac. The residue was reconstituted in 50% methanol, centrifuged again (17,000 × *g*, 10 min), and subjected to LC-MS for arginine analysis. For pyruvate derivatization, the dried extracts (prepared as above) were reconstituted in 50 µL of water and mixed with 50 µL of a derivatization buffer containing 0.5 M EDC and 0.25 M OBHA. Both EDC and OBHA suspensions were prepared in pyridine buffer (pH 5.0). The reaction mixture was extracted with 400 µL of ethyl acetate, centrifuged at 1,200 × *g* for 10 min, and the supernatant was dried using a SpeedVac. The dried residue was reconstituted in 50% methanol and centrifuged (17,000 × *g*, 10 min) prior to LC-MS analysis.

#### LC-MS/MS quantification of arginine and pyruvate

Arginine was quantified using an ExionLC™ AD series UHPLC system coupled to a QTRAP® 6500+ mass spectrometer (Sciex, Framingham, MA, USA). Separation was performed on an ACQUITY UPLC BEH Amide column (100 × 2.1 mm, 1.7 µm; Waters). Pyruvate quantification was performed on an Agilent 1290 Infinity II UHPLC system coupled to an Agilent 6460 Triple Quadrupole Mass Spectrometer (Agilent Technologies, Santa Clara, CA, USA). Separation was achieved using an ACQUITY UPLC HSS T3 column (100 × 2.1 mm, 1.8 µm; Waters) maintained at 25 °C. Detailed chromatographic conditions and MS parameters are provided in **Supplementary Methods.**

### Untargeted metabolomics analysis

#### Extraction of intra- and extracellular metabolites

*G. uro* was cultured in 50 mL PYG+ or PBM and supplemented with EA as described above. Cultures were centrifuged (8,500 × g, 5 min) to separate the supernatant (extracellular fraction) from the cell pellet (intracellular fraction). For intracellular extraction, cell pellets were resuspended in 600 µL of ice-cold methanol and flash-frozen in liquid nitrogen. After thawing, 150 µL of chloroform was added, and the samples were frozen in liquid nitrogen. Following the second thaw, 450 µL of water was added, and the mixture was vortexed and centrifuged (14,000 × *g*, 1 min). The upper aqueous layer was collected, centrifuged (17,000 × *g*, 10 min) to remove particulates, and dried using a SpeedVac. Residues were reconstituted in 80% methanol containing 1% formic acid.

For extracellular extraction, 500 µL culture supernatant was mixed with 600 µL pure methanol and centrifuged at 18,000 × *g* for 20 min. The supernatant was dried using a SpeedVac and reconstituted in 100 µL of 80% methanol containing 0.1% formic acid. For both fractions, 4-chloro-phenylalanine (400 ng/mL) was added as an IS. Intracellular metabolite data were normalized to total protein content determined by the BCA assay.

#### UHPLC-Orbitrap/MS instrumentation and analysis

Untargeted metabolic profiling was performed using a Dionex UltiMate 3000 UHPLC system coupled to an Orbitrap IQ-X Tribrid mass spectrometer (Thermo Fisher Scientific, Waltham, MA, USA). Chromatographic separation was performed on an ACQUITY UPLC BEH Amide column (100 × 2.1 mm, 1.7 µm; Waters). Detailed chromatographic conditions and LC-MS parameters are provided in **Supplementary Methods.**

### Isotope tracer analysis

#### 13C-isotope tracing of pyruvate metabolism in *G. uro* and sample preparation

*G. uro* was cultured in 2 mL PBM with isotope-labelled pyruvate (1 g/L) and supplemented with EA followed by 48-h anaerobic incubation at 37 °C. For metabolite extraction, 1 mL of bacterial cultures were collected and immediately centrifuged at 12,000 × g for 5 min at 4°C. The supernatant was collected as the extracellular fraction, while the remaining cell pellet was processed as the intracellular fraction. For intracellular extracts, the cell pellets were immediately quenched by adding 600 µL of ice-cold 80% methanol spiked with succinate-d_4_ as IS at a final concentration of 1 µM, followed by a liquid nitrogen freeze-thaw cycle. The cell pellets were further lysed by adding 150 µL of chloroform and 450 µL of deionized water, followed by another liquid nitrogen freeze-thaw step. For extracellular extracts, 500 µL of the extracellular fraction was extracted with 600 µL of ice-cold 80% methanol spiked with 1 µM succinate-d_4_ and 150 µL of chloroform. Both intracellular and extracellular extracts were then centrifuged at 17,000 × g for 10 min. The upper aqueous layer containing the intracellular metabolites was collected and vacuum-dried using Speedvac. Derivatization of pyruvate, acetate, lactate, butyrate, glutamate, and tricarboxylic acid (TCA) cycle metabolites was performed as described above.

#### LC-MS/MS instrumentation and analysis of isotope-labelled metabolites

Isotopic labeling patterns of the metabolites were evaluated using UHPLC-Orbitrap-IQX/MS coupled to a Dionex UltiMate 3000 UHPLC system (Thermo Scientific, Waltham, MA, USA). For the detection of TCA cycle intermediates (citrate, succinate, malate, fumarate, OAA, aKG) and carboxylic acids involved in the metabolic pathway of pyruvate (pyruvate, butyrate, glutamate, acetate and lactate), chromatographic separation of these OBHA-derivatized metabolites was achieved using an ACQUITY UPLC HSS T3 (2.1 mm × 100 mm, 1.8 µm, Waters Corporation). For detection of acetyl-CoA, due to its poor retention on reverse phase column, an ACQUITY UPLC HILIC column (2.1 mm × 100 mm, 1.7 µm, Waters Corporation) was used for chromatographic separation ^32^. Detailed chromatographic conditions and MS parameters are provided in **Supplementary Methods.**

The data was processed by TraceFinder software 5.1 (Thermo Scientific, Waltham, MA, USA). The metabolites of interest, including both OBHA-derivatized organic acids and non-derivatized acetyl-CoA, were identified using a combination of accurate mass (within a 5-ppm mass tolerance window) and retention time matching against authentic chemical standards. To ensure precise identification of all isotopic variants, the theoretical m/z values for each mass isotopologue were calculated using the “enviPat” isotope pattern tool. Following extraction and integration of isotopologue peaks in TraceFinder, the raw intensities were subjected to natural isotope abundance correction using the R package “AccuCor” (version 0.2.4; https://github.com/XiaoyangSu/AccuCor), which utilized the theoretical isotope distributions to generate a corrected Mass Isotopomer Distribution (MID) table. The final data were expressed as fractions.

### RNA extraction and sequencing

*G. uro* was cultured in 10 mL of PYG+, PYG+(+pyr), PBM, and PYG+(-arg), five biological replicates per condition, each supplemented with 40 µM EA. After 48 hours of anaerobic incubation, cells were harvested by centrifugation (8,500 × g, 5 min, 4 °C). The cell pellets were washed once with ice-cold PBS, snap-frozen in liquid nitrogen, and stored at -80 °C until extraction. Total RNA was extracted using a modified TRIzol-based bead-beating protocol (see **Supplementary Methods** for full details). The quality and concentration of the extracted RNA were verified using a NanoDrop One spectrophotometer (Thermo Fisher Scientific, Waltham, MA, USA). RNA sequencing (RNA-seq) library construction, sequencing, and data analysis were performed by LC-Bio Technology Co., Ltd. (Hangzhou, China) as described in **Supplementary Methods**.

### RT-qPCR analysis of hydrogen and pyruvate metabolism-related genes in *G. uro*

To validate the RNA-seq data, the expression levels of key genes involved in hydrogen and pyruvate metabolism were quantified using RT-qPCR. Total RNA was extracted using the TRIzol reagent and was reverse transcribed using the HiScript IV All-in-One Ultra RT SuperMix (Vazyme, Nanjing, China). Candidate genes, identified as significantly differentially expressed in the RNA-seq analysis, were cross-referenced with the NCBI database for primer design. Specific primers were designed using Primer-BLAST ^33^ and are listed in **Supplementary Table 1**. The cDNA was then mixed with ChamQ Universal SYBR qPCR Master Mix (Vazyme, Nanjing, China) and specific primers, followed by real-time qPCR analysis using the QuantStudio 5 Flex Real-Time PCR System (Applied Biosystems, MA, USA). The gene expression levels were normalized to *ffh* and *gyrB* and expressed as relative mRNA expression.

### Quantification of hydrogen production by gut bacteria

#### Hydrogen production by *G. uro*

To quantify hydrogen production, *G. uro* was cultured in 200-mL serum bottles sealed with butyl rubber stoppers and aluminum crimp caps to prevent gas exchange. Each bottle contained 170 mL of PYG+ or PBM, leaving a 30-mL headspace for gas accumulation. The medium-filled bottles were sterilized at 121 °C for 20 min. To ensure anaerobic conditions prior to inoculation, the bottles were placed in an anaerobic jar containing gas-generating sachets with the caps open for 24 h. Following this pre-reduction step, the media were inoculated (1:100, v/v) with *G. uro*, and the bottles were immediately sealed tightly with butyl rubber stoppers and aluminum crimp caps. The cultures were incubated at 37 °C for 72 h, after which gas samples were collected from the headspace for GC-TCD analysis.

### Hydrogen production by *Bacteroides* spp

The hydrogen-producing capacity of *B. thetaiotaomicron*, *B. vulgatus*, and *B. stercoris* was evaluated using the same protocol mentioned above. Strains were cultured in 170 mL of WCM in serum bottles, pre-reduced with open caps as described above, sealed, and incubated at 37 °C for 48 h prior to headspace analysis.

#### GC-TCD instrumentation and quantification of hydrogen levels

Headspace gas samples were collected using a gastight syringe and analyzed via gas chromatography equipped with a thermal conductivity detector (GC-TCD; Agilent 8890). Separation was achieved using a HayeSep Q column (3 ft × 1/8 in × 2 mm, 80/100 mesh; Agilent G3591-81020) and MolSieve 5A column (8 ft × 1/8 in × 2 mm, 60/80 mesh; Agilent G3591-81022). Nitrogen gas was used as the carrier gas, the flow rate for the HayeSep Q column was maintained at 26 mL/min, while the Molecular Sieve 5A column was operated at a constant forward pressure of 25 psi. The injector and detector temperatures were 250 °C, and the initial column temperature was set at 60 °C for 1 min, ramped at 20 °C/min to 80 °C, then at 30 °C/min to 190 °C held for 0.3 min.

### Effects of hydrogen exposure on EA-to-urolithin conversion

#### EA-to-UroC conversion by *G. uro* with hydrogen supplementation

To investigate the direct influence of exogenous hydrogen on urolithin production, *G. uro* was cultured in the presence of a commercial hydrogen-producing supplement (H₂ Molecular Hydrogen, Dr. Mercola, FL, USA). Each tablet is formulated to generate approximately 8 ppm of molecular hydrogen. *G. uro* was inoculated into 170 mL of PYG+ or PBM within the serum bottles as described above. Immediately prior to sealing, one hydrogen tablet was added to each bottle to initiate hydrogen release. The bottles were promptly capped with butyl rubber stoppers and aluminum seals to retain the generated gas. Cultures were supplemented with 40 µM EA and incubated at 37 °C for 72 h. Following incubation, aliquots were collected for metabolite extraction and LC-MS analysis of EA and urolithins.

#### EA-to-UroC conversion by *G. uro* co-cultured with *Bacteroides*

To evaluate the synergistic effect of hydrogen-producing bacteria on EA biotransformation, *G. uro* was co-cultured with selected *Bacteroides* strains. Assays were performed in 96-well plates containing 250 µL of PYG+ or PBM. The media were supplemented with 40 µM EA and inoculated with *G. uro* (1:100 dilution) alongside one of the following strains (*B. thetaiotaomicron*, *B. vulgatus*, or *B. stercoris*) at an equivalent inoculation ratio. Monocultures of *G. uro* and each *Bacteroides* strains alone served as controls. Following anaerobic incubation at 37 °C for 72 h, samples were processed for EA and urolithin extraction and analyzed via LC-MS as described above.

### Assessment of EA-to-urolithin conversion *in vivo*

To investigate the functionality of the optimized PBM conditions in a gut microbiota environment, we conducted an animal experiment assessing EA-to-urolithin conversion efficiency. Male C57BL/6J mice (5-6 weeks old) were obtained from the centralized animal facilities at The Hong Kong Polytechnic University. Mice were housed under standard conditions with a 12-h light/dark cycle (07:30–19:30), a temperature of 21 ± 1°C, and humidity of 65% ± 5%. All animal procedures were approved by the Hong Kong Polytechnic University Animal Subjects Ethics Sub-committee (ASESC project number: 23-24/901-FSN-R-GRF).

After two weeks of acclimation, 12 mice were randomly assigned to two groups (n = 6 per group). To minimize background arginine intake, the standard chow diet was replaced with a low-arginine diet consisting exclusively of boiled potato for the duration of the experiment. Mice were transferred to metabolic cages 12 hours prior to substrate administration (-12 h) and maintained there until the study endpoint (+60 h) to facilitate sample collection. The experimental groups differed by their fluid intake and specific gavage treatments: the PBM group received PBM medium supplemented with 3 g/L sodium pyruvate as their sole drinking source, along with a single oral gavage of sodium pyruvate (1 g/kg body weight) at 0 h; the PYG+ group received PYG+ medium containing 15 g/L arginine as their drinking source, along with a single oral gavage of L-arginine (3 g/kg body weight) at 0 h.

To ensure consistent bacterial colonization, all mice were orally administered fresh *G. uro* cultures (∼9.0 log CFU/mouse) at three time points: -48 h, -24 h, and +18 h relative to substrate administration. EA (30 mg/kg body weight), suspended in 0.4% CMC-Na, was administered as a single oral dose at 0 h. Urine and fecal samples were collected at 0, 18, 38, and 60 h post-administration and immediately stored at -80°C until metabolite analysis. Extraction and quantification of EA and urolithins were performed as described above.

### Statistical analysis

All statistical tests were performed with the GraphPad Prism software (v.10.1.2). Normal distributions were evaluated by the Shapiro–Wilk test. For multiple groups comparison, statistical significance was analysed by one-way ANOVA, followed by Tukey’s post-hoc test while for comparison between two groups, unpaired two-tailed Student’s t-test was used. Data were plotted in GraphPad Prism (v10.0.0) and R programming (v.4.3.2).

## Supporting information

PBM Manuscript-17 Apr-CL.pdf

## ACKNOWLEDGEMENT

The authors wish to acknowledge the Innovation and Technology Fund-Innovation and Technology Support Programme (ITF-ITSP) of the Innovation and Technology Commission of Hong Kong (ITS/040/22MS), the General Research Fund-Early Career Scheme (25300323) and the Young Collaborative Research Fund (C4004-23Y) from the Research Grants Council (RGC) of Hong Kong for the funding. We also greatly appreciate technical support from the University Research Facility in Life Science (ULS) and the University Research Facility in Chemical and Environmental Analysis (UCEA) at the Hong Kong Polytechnic University.

## AUTHOR CONTRIBUTIONS

M.E.G. and D.Z. conceived and designed the research. M.E.G. performed the majority of the experiments, data analysis, and wrote the original draft of the manuscript. P.K.L. conducted the ¹³C-isotopic tracer experiment, assisted in data analysis, and drafted parts the manuscript. J.L. conducted the untargeted metabolomics experiments and data analysis. L.Z. and X.X. conducted bacterial genetic engineering in *G. uro*. M.Z. and M.J.L. assisted in the hydrogen measurement experiments. X.Y. assisted in fermentation experiments for assessing polyphenol conversions and data analysis. L.D. provided technical oversight, essential resources, and revised the manuscript. D.Z. provided the funding and essential resources, and directed the overall research. All authors contributed to the discussion of the results, revision of the manuscript, and approved the final version of the manuscript.

