## Supplementary material for "Pyruvate-driven hydrogen production promotes polyphenol bioconversion by gut bacteria": PBM Manuscript-17 Apr-CL.pdf

***Supplementary Methods***

**Effect of hemin concentration on ellagic acid bioconversion.** To assess the impact of hemin on EA metabolism, *G. uro* was cultured in modified PBM containing varying concentrations of hemin: 0 mg/L (hemin-free), 5 mg/L (standard PBM), and 10 mg/L. The cultures were supplemented with EA and incubated anaerobically in 96-well plates at 37 °C for 3 days. Following incubation, the conversion efficiencies of EA to UroC and total urolithins were quantified using LC-MS/MS, as described in the main methodology.

**LC-MS/MS quantification of phenolic metabolites**. LC-MS/MS analysis was performed using an ExionLC™ AD series UHPLC system coupled to a QTRAP® 6500+ mass spectrometer (Sciex, Framingham, MA, USA) equipped with an electrospray ionization (ESI) source. Chromatographic separation was achieved using an ACQUITY UPLC HSS T3 column (100 × 2.1 mm, 1.8 μm; Waters, Milford, MA, USA) maintained at 40 °C. The mobile phases consisted of water containing 0.1% formic acid (A) and acetonitrile containing 0.1% formic acid (B). The gradient elution, denoted as B%, starting at 5% from 0 to 1 min, increasing to 65% at 10 min, reaching 99% from 12 to 14 min, and then returning to the initial condition in 0.1 minute. The flow rate was 0.4 mL/min and the sample injection volume was 8 μL. The targeted phenolic metabolites were detected in negative ion mode with the following optimized source parameters: ion spray voltage: -4500 V; temperature: 550 °C; ion source gas 1: 60 psi; and ion source gas 2: 60 psi. The MS/MS parameters of target metabolites and internal standards are listed in Supplementary Table 4. Data acquisition and processing were performed using Analyst software (version 1.7.2, Sciex).

**LC-MS/MS quantification of arginine and pyruvate.** Arginine was quantified using an ExionLC™ AD series UHPLC system coupled to a QTRAP® 6500+ mass spectrometer (Sciex, Framingham, MA, USA). Chromatographic separation was performed on an ACQUITY UPLC BEH Amide column (100 × 2.1 mm, 1.7 μm; Waters). The mobile phases consisted of water containing 0.1% formic acid (A) and acetonitrile containing 0.1% formic acid (B). The gradient elution, denoted as B%, starting at 1% from 0 to 0.5 min, increasing to 50% at 7 min, reaching 95% from 10 to 13 min, and then returning to the initial condition in 0.1 minute. The flow rate was 0.3 mL/min and the sample injection volume was 2 μL. Pyruvate quantification was performed on an Agilent 1290 Infinity II UHPLC system coupled to an Agilent 6460 Triple Quadrupole Mass Spectrometer (Agilent Technologies, Santa Clara, CA, USA). Chromatographic separation was achieved using an ACQUITY UPLC HSS T3 column (100 × 2.1 mm, 1.8 μm; Waters) maintained at 25 °C. The mobile phases consisted of water containing 0.1% formic acid (A) and acetonitrile containing 0.1% formic acid (B). The gradient elution, denoted as B%, starting at 2% from 0 to 2 min, reaching 60% from 4.5 to 9 min, increasing to 80% at 10 min, and then returning to the initial condition in 0.5 minute. The flow rate was 0.3 mL/min and the sample injection volume was 5 μL. Both arginine and pyruvate were detected in positive ion mode with the following optimized source parameters: ion spray voltage: 5000 V; temperature: 600 °C; ion source gas 1: 60 psi; and ion source gas 2: 60 psi. The MS/MS parameters of target metabolites and internal standards are listed in Supplementary Table 5.

**Untargeted metabolomics analysis**. Untargeted metabolic profiling was performed using a Dionex UltiMate 3000 UHPLC system coupled to an Orbitrap IQ-X Tribrid mass spectrometer (Thermo Fisher Scientific, Waltham, MA, USA). Chromatographic separation was performed on an ACQUITY UPLC BEH Amide column (100 × 2.1 mm, 1.7 µm; Waters). The mobile phases consisted of Milli-Q water containing 5 mM ammonium formate and 0.1% formic acid (A) and acetonitrile:Milli-Q water (95:5, v/v) containing 5 mM ammonium formate and 0.1% formic acid (B). The gradient elution, denoted as B%, starting at 98% from 0 to 1 min, decreasing to 70% at 8 min, reaching 50% from 10 to 12 min, and then returning to the initial condition in 0.1 minute. The flow rate was 0.3 mL/min and the sample injection volume was 3 µL. Data acquisition was operated in full scan ESI mode with a scan range of 70–1000 m/z. The ESI source parameters were set as follows: Drying gas temperature: 300 °C; Drying gas flow: 8 L/min; Nebulizer pressure: 40 psi; Sheath gas temperature: 320 °C; Sheath gas flow: 11 L/min; Capillary voltage (VCap): 4000 V; Fragmentor voltage: 140 V; Skimmer voltage: 65 V.

**LC-MS/MS instrumentation and analysis of isotope-labelled metabolites**. For the detection of TCA cycle intermediates (citrate, succinate, malate, fumarate, OAA, aKG) and carboxylic acids involved in the metabolic pathway of pyruvate (pyruvate, butyrate, glutamate, acetate and lactate), chromatographic separation was achieved using an ACQUITY UPLC HSS T3 (2.1 mm × 100 mm, 1.8 µm, Waters Corporation), maintained at 40 °C. The mobile phases consisted of water containing 0.1% formic acid (A) and acetonitrile containing 0.1% formic acid (B). The gradient elution, denoted as B%, starting at 2% from 0 to 2 min, increasing to 60% at 5 min, reaching 90% from 7 to 9 min, and then returning to the initial condition in 0.1 minute. For detection of acetyl-CoA, due to its poor retention on reverse phase column, an ACQUITY UPLC HILIC column (2.1 mm × 100 mm, 1.7 µm, Waters Corporation), maintained at 40 °C, was used for chromatographic separation with reference to Singh et al. (2024). The mobile phases consisted of acetonitrile:Milli-Q water (1:9, v/v) with 5 mM ammonium acetate (A) and acetonitrile:Milli-Q water (9:1, v/v) with 5 mM ammonium acetate (B). The gradient elution, denoted as B%, starting at 90% from 0 to 0.5 min, decreasing to 50% at 5 min, reaching 50% from 5 to 8 min, and then returning to the initial condition in 0.1 minute. The flow rate was 0.3 mL/min and the sample injection volume was 3 µL. Data acquisition was operated in full scan positive ESI mode with a scan range of 70–3000 m/z. The ESI source parameters were set as follows: Drying gas temperature: 300 °C; Drying gas flow: 8 L/min; Nebulizer pressure: 40 psi; Sheath gas temperature: 320 °C; Sheath gas flow: 11 L/min; Capillary voltage (VCap): 4000 V; Fragmentor voltage: 140V; Skimmer voltage: 65 V.

**RNA extraction and quality assessment**. Total RNA was extracted using the TRIzol reagent (Takara Bio, Kusatsu, Japan) with a modified bead-beating protocol. Briefly, frozen pellets were resuspended in 1.0 mL of TRIzol and transferred to screw-cap tubes containing 4-5 small glass beads. Cell lysis was performed using a homogenizer (Precellys Evolution, Bertin Technologies, France) set to level 3 for 2–3 cycles of 30 s each. Following lysis, 0.2 mL of chloroform was added, and centrifuged at 12,000 × g for 15 min at 4 °C. The upper aqueous phase was transferred to a new tube, mixed with 0.5 mL of isopropanol, and incubated at room temperature for 10 min to precipitate the RNA. The RNA was pelleted by centrifugation at 12,000 × g for 15 min at 4 °C. The supernatant was discarded, and the pellet was washed with 1 mL of 75% ethanol followed by centrifugation at 12,000 × g for 5 min at 4 °C. The washed pellet was air-dried for 10 min, resuspended in 30 μL of DEPC-treated water, and incubated at 55 °C for 10 min to ensure solubilization. RNA purity was assessed using a NanoDrop spectrophotometer, and concentration was precisely quantified using a Qubit fluorometer. RNA integrity was evaluated by 1% agarose gel electrophoresis. Samples with clear ribosomal RNA bands (23S/16S) and RNA integrity suitable for library construction were used for subsequent steps.

**RNA-seq library construction, sequencing, and data analysis.** For each sample, ribosomal RNA was depleted using appropriate probes, followed by DNase I treatment to eliminate potential DNA contamination. RNA was then fragmented into short pieces using fragmentation buffer. First-strand cDNA was synthesized using random hexamer primers, followed by second-strand cDNA synthesis. The double-stranded cDNA was purified, subjected to end repair, A-tailing, and ligation of sequencing adapters. After fragment size selection using AMPure XP beads, the final cDNA library was obtained by PCR amplification. Library quality was assessed by quantifying concentration and verifying insert size. Qualified libraries were sequenced by LC-Bio Technology Co., Ltd. (Hangzhou, China) on an Illumina platform (paired-end 150 bp reads) according to the manufacturer's instructions. Sequencing was conducted in two independent batches: the first batch included the PBM and standard PYG+ groups, while the second batch included the PBM, PYG+(-arg), and PYG+(+pyr) groups. Raw sequencing reads were processed using fastp (version 0.21.0) to remove adapter sequences and low-quality reads. Clean reads were mapped to the *G. uro* reference genome using Bowtie2 (version 2.4.5). Gene expression levels were quantified as fragments per kilobase per million mapped reads (FPKM) using HTSeq-count. Differential expression analysis between experimental groups was performed using DESeq2 (version 1.26.0) with thresholds of $|\log_2 FoldChange| > 1$ and adjusted P value $< 0.05$. Gene Ontology (GO) enrichment and KEGG pathway analyses were conducted using cluster Profiler (version 3.14.3).

**Quantification of short-chain fatty acids (SCFAs).** SCFA production by *G. uro* was evaluated in PYG+, PYG+(+pyr), PBM, and PYG+(-arg) supplemented with 40 μM EA. After 72 h of anaerobic incubation at 37°C, culture samples were collected for analysis. Extraction and quantification were performed as previously described [1].

**Effect of glucose on EA bioconversion.** To determine whether the presence of glucose in PBM influences EA bioconversion, a manual, glucose-free PBM was formulated and compared against standard PBM. PBM was prepared using commercial PYG, which inherently contains 5 g/L glucose. The glucose-free PBM was prepared using a manually compounded PYG base consisting of 5 g/L tryptone, 5 g/L peptone, 10 g/L yeast extract, 5 g/L beef extract, 2 g/L K2HPO4, 1 mL/L Tween 80, 0.5 g/L L-cysteine-HCl·H2O, and 40 mL/L of a salt solution (0.25 g/L CaCl2·2H2O, 0.5 g/L MgSO4·7H2O, 1 g/L K2HPO4, 1 g/L KH2PO4, 10 g/L NaHCO3, and 2 g/L NaCl). Both PBM and glucose-free PBM were then supplemented with 2 g/L dipotassium phosphate, 1 mL/L Tween 80, 1 g/L pyruvate, 5 mg/L hemin, 1 mg/L vitamin K, and 1 mg/L resazurin. *G. uro* cultures were supplemented with EA and incubated anaerobically in 96-well plates at 37 °C for 3 days. EA conversion efficiency was subsequently evaluated using LC-MS/MS.

**Measurement of dissolved hydrogen production using methylene blue-based hydrogen titration.** Dissolved hydrogen accumulation in the culture media was quantified using a methylene blue-platinum titration reagent (H2Blue; Kueysing, China). Briefly, *G. uro* was cultured anaerobically in 175 mL of either PYG+ or PBM, both supplemented with ellagic acid (EA), in sealed serum bottles for 3 days. Following incubation, the culture broth was titrated with the H2Blue reagent according to a modified manufacturer's colorimetric drop-count method. The sample was titrated dropwise until the blue color no longer faded, with each drop required to neutralize the dissolved hydrogen corresponding to 0.1 ppm. Bacteria-free PYG+ and PBM media were incubated in parallel under identical conditions to serve as baseline controls.

**Effect of exogenous hydrogen supplementation on EA bioconversion.** To evaluate the impact of an exogenous hydrogen source on EA metabolism, *G. uro* was cultured in 175 mL of either PYG+ or PBM. Both media were supplemented with EA and prepared in sealed serum bottles (pre-incubated in anaerobic condition for 24 h) to maintain anaerobic conditions and trap generated gases. Commercial hydrogen-producing tablets ($H_2$ Molecular Hydrogen, Dr. Mercola, FL, USA) were added to the experimental cultures at doses of either 2 full tablets (high dose) or 1/2 (low dose) tablet per bottle. The cultures were incubated anaerobically at 37 °C for 3 days. Following the incubation period, the bioconversion of EA to urolithins was quantified using LC-MS/MS, consistent with the main methodology.

***Supplementary Figures***

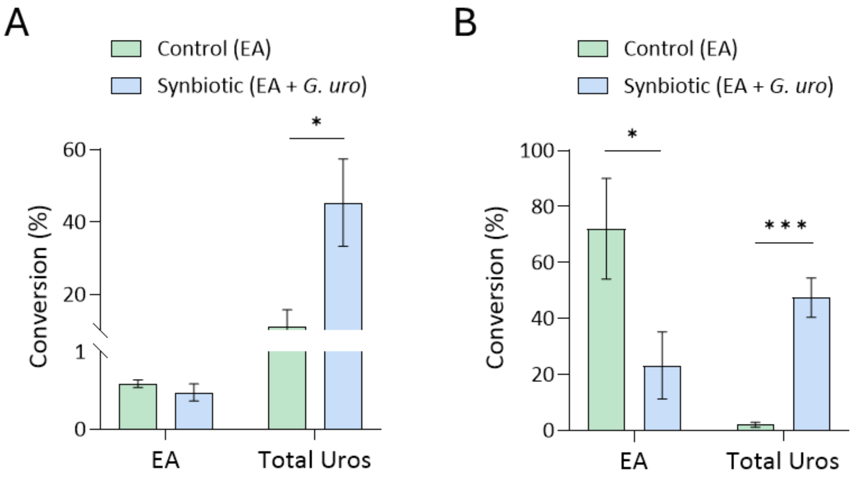


**Supplementary Fig. 1. A synbiotic combination of *G. uro* and EA efficiently promotes EA-to-urolithins conversion in mice. (A)** Urinary and **(B)** fecal excretion of ellagic acid (EA) and total Uros (including UroM7, UroM6, UroD, UroC, UroA, dimethyl UroC, methyl UroA, and methyl UroB) over 36 h in mice administered a single oral dose of 30 mg/kg EA (control) or a synbiotic combination of EA and *G. uro*. Data are presented as mean ± SEM ($n$ = 5). Statistical significance was assessed using Student's t-test (*, $P < 0.05$; **, $P < 0.01$; ***, $P < 0.001$).

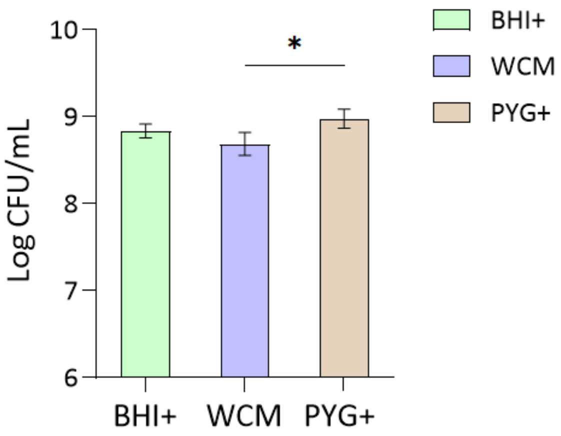


**Supplementary Fig. 2. *G. uro* growth in different standard growth media.** *G. uro* was cultured in BHI+, WCM, and PYG followed by 72 h anaerobic incubation. Data are presented as mean ± SEM ($n = 3$). Statistical significance was assessed using one-way ANOVA followed by Tukey's post hoc test (multiple groups) (*, $P < 0.05$).

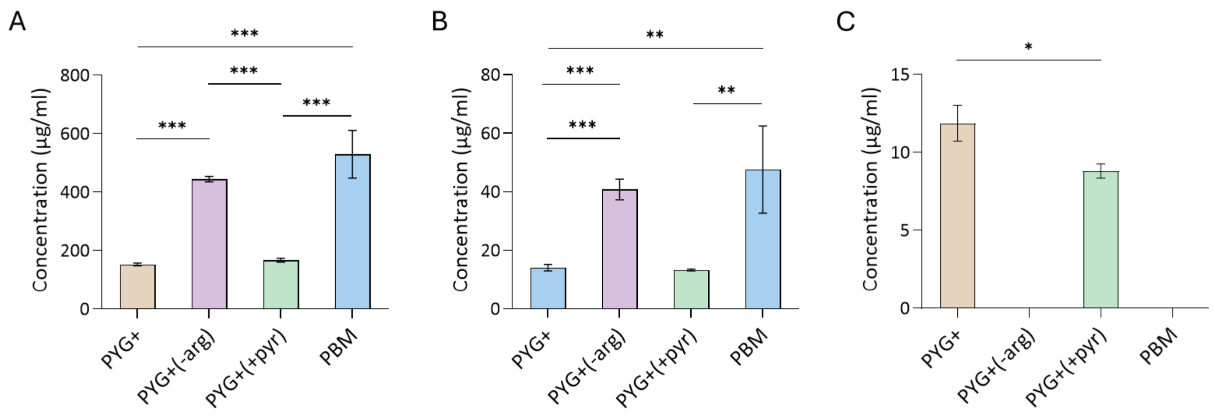


**Supplementary Fig. 3. Production of short-chain fatty acids (SCFAs) by *G. uro*. (A-C)** Extracellular concentrations of (A) acetic acid, (B) valeric acid, and (C) isovaleric acid in PYG+, PYG+(-arg), PYG+(+pyr), and PBM. Bacterial cultures were supplemented with EA (40 µM) and incubated anaerobically for 72 h. Data are presented as mean ± SEM ($n = 3$). Statistical significance was assessed using one-way ANOVA followed by Tukey's post hoc test (*, $P < 0.05$; **, $P < 0.01$; ***, $P < 0.001$).

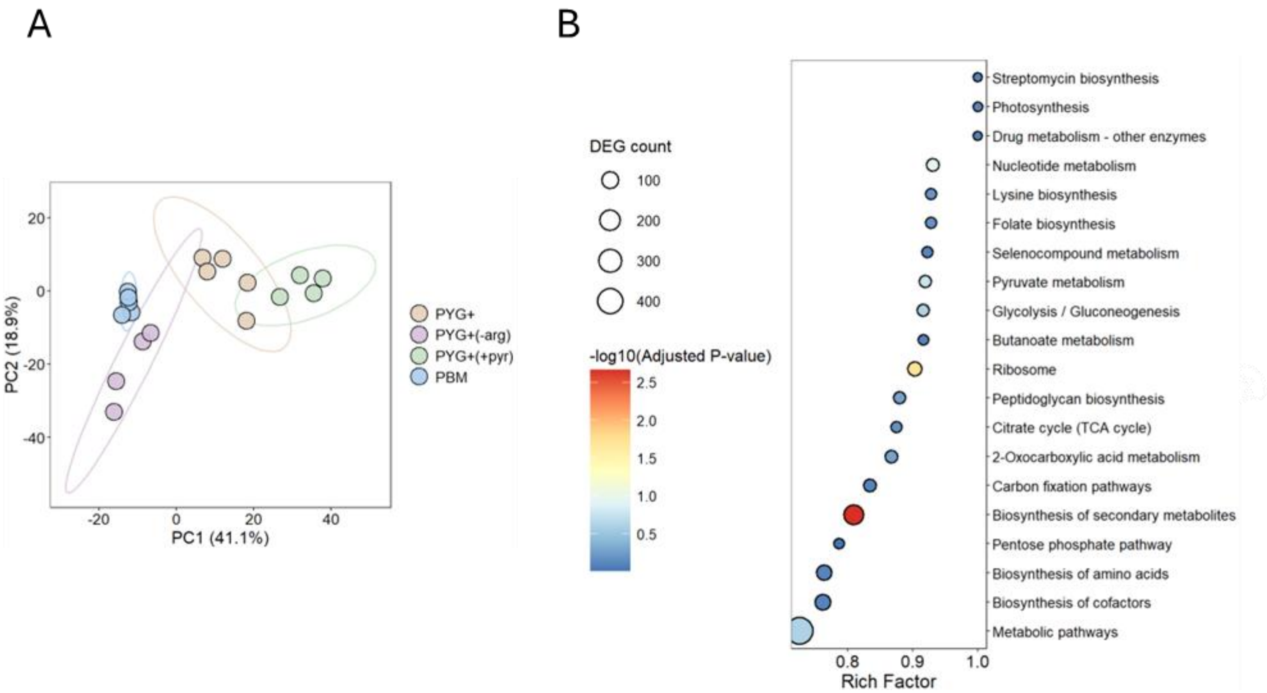


**Supplementary Fig. 4. Transcriptomic profiling of *G. uro* in different media**. **(A)** PCA of RNA-seq data from *G. uro* cultured in four media: PBM, PYG+, PYG+(-arg), and PYG+(+pyr). **(B)** Pathway enrichment analysis of DEGs significantly upregulated in PBM relative to PYG+(+pyr). Bubble size represents the DEG count, and color intensity indicates the P value. *G. uro* was cultured anaerobically in EA-supplemented PBM or PYG+ for 48 h, followed by RNA isolation and sequencing. Pathway enrichment was determined by hypergeometric test with FDR correction.

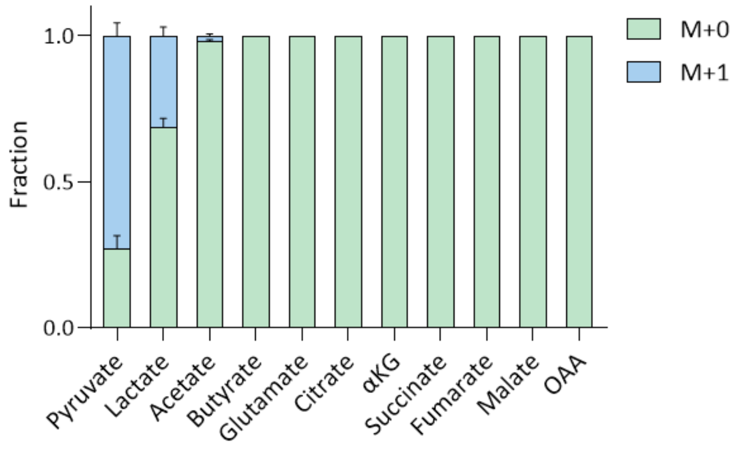


**Supplementary Fig. 5. Stable isotope tracer analysis of pyruvate metabolism in *G. uro* using [3-$^{13}$C]-pyruvate combined with LC-MS.** Mass isotopologue distributions of extracellular pyruvate-derived organic acids. *G. uro* cultures in PBM were supplemented with EA and [3-$^{13}$C]-pyruvate and incubated anaerobically for 48 h. All data are presented as mean ± SEM (n = 3). α-KG, α-ketoglutarate; OAA, oxaloacetate.

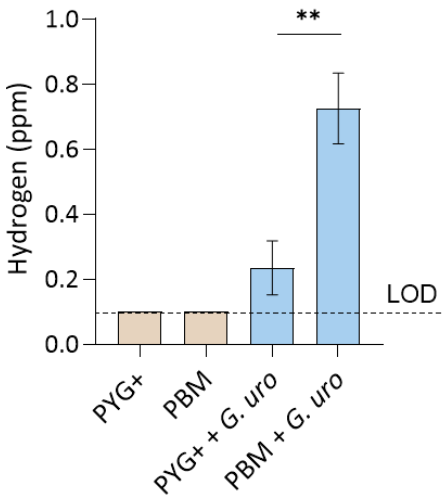


**Supplementary Fig. 6. Quantification of dissolved hydrogen production by *G. uro* in PYG+ and PBM.** Dissolved hydrogen concentration (ppm) was measured after 72 h anaerobic incubation in sealed serum bottles containing PYG+ or PBM supplemented with EA using H2Blue titration assay. The limit of detection (LOD = 0.1 ppm) is indicated by the dotted line. Data are presented as mean ± SEM ($n$ = 4). Statistical significance was assessed using Student's t-test (*, $P < 0.05$; **, $P < 0.01$; ***, $P < 0.001$).

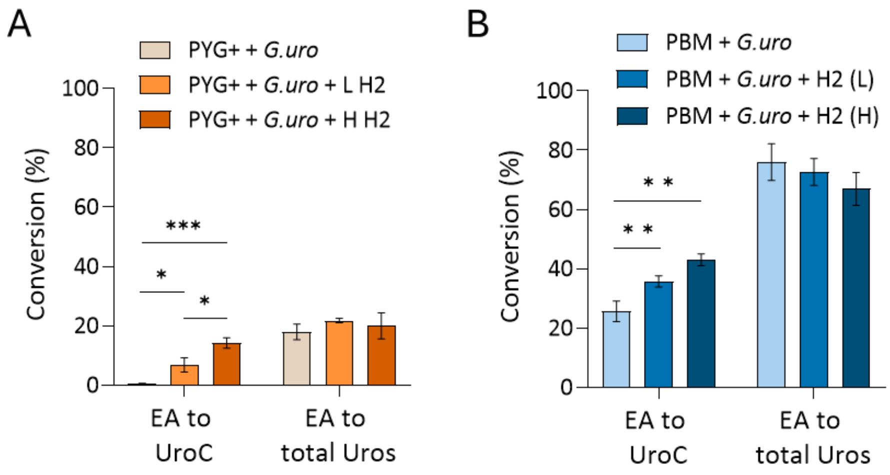


**Supplementary Fig. 7. Effect of exogenous hydrogen supplementation on EA-to-urolithin conversion by *G. uro*. (A-B)** Conversion of EA to UroC and total Uros were measured in after 72-h culturing *G. uro* in sealed serum bottles containing (A) PYG+ or (B) PBM and supplemented with high (H) and low (L) doses of hydrogen-producing tablets. Data are presented as mean ± SEM ($n$ = 3). Statistical significance was assessed using one-way ANOVA followed by Tukey's post hoc test (*, $P < 0.05$; **, $P < 0.01$; ***, $P < 0.001$).

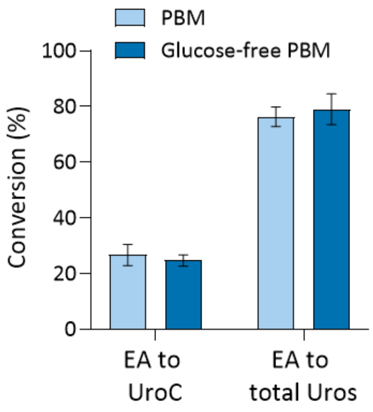


**Supplementary Fig. 8. Effect of glucose on on EA-to-urolithin conversion by *G. uro*.** Conversion of EA to UroC and total Uros in PBM (containing 5 g/L glucose) and glucose-free PBM by *G. uro*. Bacterial cultures were supplemented with EA (40 μM) and incubated anaerobically for 72 h. Data are presented as mean ± SEM ($n$ = 3). Data are presented as mean ± SEM ($n$ = 3). Statistical significance was assessed using Student's t-test (*, $P < 0.05$).

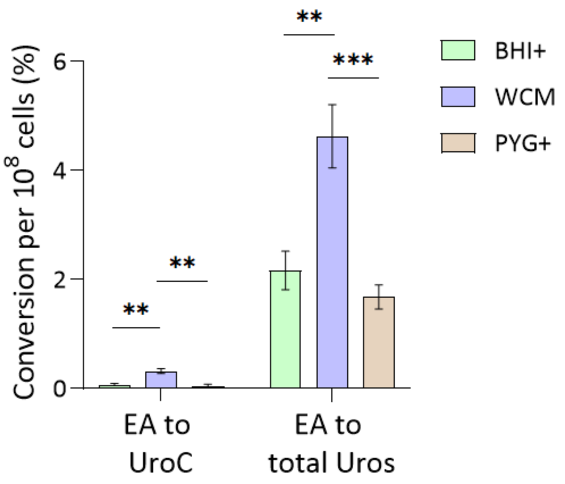


**Supplementary Fig. 9. EA-to-urolithin conversion normalized to *G. uro* cell number in standard growth media.** EA-to-urolithin conversion (Fig. 1B) and growth (Supplementary Fig. 2) data were used to normalize the conversion to bacterial counts. All data are presented as mean ± SEM ($n$ = 3). Statistical significance was assessed using one-way ANOVA followed by Tukey's post hoc test (*, $P < 0.05$; **, $P < 0.01$; ***, $P < 0.001$).

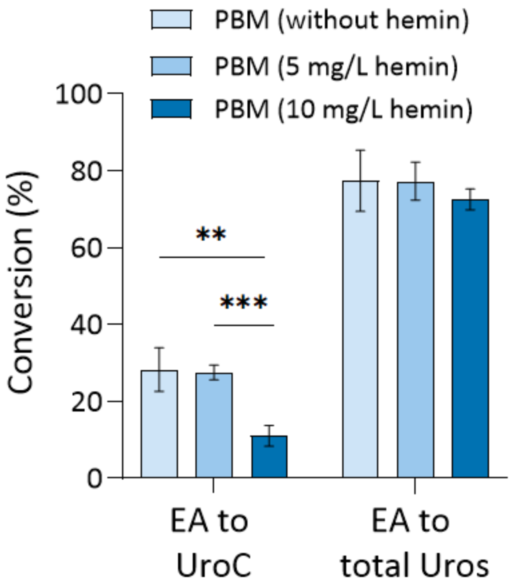


**Supplementary Fig. 10. Effect of hemin concentration on EA-to-urolithin conversion by *G. uro*.** Conversion of EA to UroC and total Uros cultured in PBM containing 0 mg/L, 5 mg/L (standard PBM), or 10 mg/L of hemin. Bacterial cultures were supplemented with EA (40 μM) and incubated anaerobically for 72 h. Data are presented as mean ± SEM (*n* = 3). Statistical significance was assessed using one-way ANOVA followed by Tukey's post hoc test (*, $P < 0.05$; **, $P < 0.01$; ***, $P < 0.001$).

*Supplementary Tables*

**Supplementary Table 1.** Primer sequences used for RT-qPCR validation.

| Gene Name | Forward Primer (5' → 3') | Reverse Primer (5' → 3') | Functions |
|---|---|---|---|
| nifJ | GTCTCCTTACGCGGTGAACA | CGGAGGCGGTGTCCTTATAC | Pyruvate:ferredoxin (flavodoxin) oxidoreductase |
| *porA* | GTCATGATGAACGCGAACCG | ACGGCATAGGCCATAAGAGC | Pyruvate ferredoxin oxidoreductase |
| *hypD* | CGATCGCGAGATCGTGTTCG | ACCGGGCATGTTCTTGTGGG | Hydrogenase formation protein HypD |
| *hypB* | GTCATGATCCTGTCCGTTCC | GAAGTTGAACACGGGCAGGG | Hydrogenase nickel incorporation protein HypB |
| *hypA* | AATGCGGGAGCTTCGCCAC | GTCGTCGGGGATATCCACCT | Hydrogenase nickel metallochaperone HypA |
| *hypE* | GACATCACCGTGGAGGAGGA | GGATCGTAGCCCAGCATCTC | Hydrogenase expression/formation protein HypE |
| Hyd-S | GCATGAAGATCACGGGCCG | GGTGGTTCTTGTCCCAGGC | Hydrogenase small subunit |
| Hyd-L | CCGAGTACATGGGCCACTCC | TGTAGCGGTCGTTCACGTCG | Nickel-dependent hydrogenase large subunit |
| *Ffh* | TGGACTTCGACGGCGTTATC | GCCGGACGAGATGAACTTGA | Housekeeping genes |
| *gyrB* | ACTGCGACGAGATCAAGGTG | TCTTTCGGGTGCTTGTCGAT | Housekeeping genes |

**Supplementary Table 2.** Major differences in key nutrient compositions among PYG+, WCM, BHI+, and PBM broths.

| Nutrient | PYG+ | WCM | BHI+ | PBM |
|---|---|---|---|---|
| Arginine | High | Low | High | None |
| Pyruvate | Without | With | Without | With |
| Glucose | High | Without | Low | Low |
| Hemin | With | With | Without | With |

**Supplementary Table 3.** Differential expression of selected genes involved in pyruvate metabolism and hydrogenase formation in *G. uro*. Data were derived from RNA-seq analysis ($n$ = 5). Differentially expressed genes (DEGs) were identified using FDR threshold of < 0.05 with Benjamini–Hochberg correction for multiple comparisons.

| PBM relative to PYG+ | | | | | |
|---|---|---|---|---|---|
| **Pathway** | **Gene ID** | **Gene name** | **Function** | **Log2 FC PBM vs. PYG+** | **FDR** |
| Pyruvate metabolism | BN3560_ RS02785 | nifJ | pyruvate:ferredoxin (flavodoxin) oxidoreductase | 4.8 | 1.16E-20 |
| Pyruvate metabolism | BN3560_ RS05870 | porA | pyruvate ferredoxin oxidoreductase | 2.3 | 9.60E-03 |
| Hydrogenase | BN3560_ RS06430 | hypD | hydrogenase formation protein HypD | 3.0 | 6.94E-07 |
| Hydrogenase | BN3560_ RS06435 | - | HypC/HybG/HupF family hydrogenase formation | 2.6 | 1.78E-04 |
| Hydrogenase | BN3560_ RS07210 | hypE | hydrogenase expression/formation protein HypE | 3.2 | 1.47E-07 |
| Hydrogenase | BN3560_ RS08920 | - | hydrogenase maturation nickel metallochaperone HypA | 2.6 | 1.03E-04 |
| Hydrogenase | BN3560_ RS08925 | hypB | hydrogenase nickel incorporation protein HypB | 3.3 | 3.14E-09 |
| Hydrogenase | BN3560_ RS12445 | - | Ni/Fe hydrogenase subunit alpha | 5.4 | 2.13E-21 |
| Hydrogenase | BN3560_ RS14075 | - | nickel-dependent hydrogenase large subunit | 5.4 | 1.22E-21 |
| **PBM relative to PYG+(+pyr)** | | | | | |
| Pyruvate metabolism | DMP12_R S08390 | nifJ | pyruvate:ferredoxin (flavodoxin) oxidoreductase | 5.8 | 7.68E-198 |
| Pyruvate metabolism | DMP12_R S01615 | porA | pyruvate ferredoxin oxidoreductase | 2.9 | 1.16E-04 |
| Hydrogenase | DMP12_R S01055 | hypD | hydrogenase formation protein HypD | 3.5 | 2.97E-15 |
| Hydrogenase | DMP12_R S00265 | hypE | hydrogenase expression/formation protein HypE | 2.4 | 3.13E-04 |
| Hydrogenase | DMP12_R S02300 | hypB | hydrogenase nickel incorporation protein HypB | 5.0 | 9.13E-36 |

| | | | | | |
|---|---|---|---|---|---|
| Hydrogenase | DMP12_RS10905 | - | ferredoxin hydrogenase | 7.1 | 3.77E-20 |
| Hydrogenase | DMP12_RS10790 | - | ferredoxin hydrogenase | 7.9 | 7.68E-20 |

**Supplementary Table 4.** Optimized MRM parameters for targeted polyphenol analysis in negative ESI mode.

| Compound | MRM transitions<br>Precursor → Quantifier (Qualifier) ions* | DP (V) | CE (V) |
|---|---|---|---|
| Ellagic acid | 301.0 → 284.1 (145.1) | -150 | -41 |
| Urolithin C | 243.0 → 187.1 (171.0) | -150 | -40 |
| Urolithin D | 259.0 → 242.0 (213.0) | -100 | -20 |
| Urolithin M6 | 259.0 → 213.0 (159.0) | -120 | -35 |
| Urolithin M7 | 243.0 → 198.0 (147.0) | -130 | -35 |
| Urolithin B | 211.1 → 167.1 (139.1) | -110 | -33 |
| 6,7-Dihydroxycoumarin (IS) | 177.0 → 133.0 (105.1) | -85 | -25 |

*Transition from precursor ion to quantifier ion was used for quantification, and the qualifier ion was used for confirmation.

**Supplementary Table 5.** Optimized MRM parameters for targeted arginine and pyruvate analysis in positive mode.

| **Compound** | **MRM transitions**<br>**Precursor → Quantifier (Qualifier) ions*** | **DP (V)** | **CE (V)** |
|---|---|---|---|
| Arginine | 175.0 → 70.0 (128.8) | 170 | 33 |
| Pyruvate | 299.3 → 181.2 | 100 | 6 |

*Transition from precursor ion to quantifier ion was used for quantification, and the qualifier ion was used for confirmation.
